# muSim: A goal-driven framework for elucidating the neural control of movement through musculoskeletal modeling

**DOI:** 10.1101/2024.02.02.578628

**Authors:** Muhammad Noman Almani, John Lazzari, Shreya Saxena

## Abstract

How does the motor cortex (MC) produce purposeful and generalizable movements with the complex musculoskeletal system in a dynamic environment? To elucidate the underlying neural dynamics, we use a goal-driven approach to model MC by considering its goal as a controller driving the musculoskeletal system through desired states to achieve movement. Specifically, we formulate a model of MC as a recurrent neural network (RNN) controller producing muscle commands while receiving sensory feedback from biologically accurate musculoskeletal models. Given this real-time simulated feedback implemented in advanced physics simulation engines, we use deep reinforcement learning to train the RNN to execute desired movements under specified neural and musculoskeletal constraints. For general use, we provide a modular computational framework that allows the flexible integration of user-defined musculoskeletal models, training algorithms, tasks and constraints. We also provide a combination of modules to analyze and quantify the dynamical alignment and similarity of the trained RNN with the recorded neural data on the population and single-unit level. Using these modules, we find that the activity of the trained RNN can accurately decode experimentally recorded neural population dynamics and single-unit MC activity, while generalizing well to testing conditions significantly different from training. Finally, we also provide perturbation modules to generate insights about neural dynamics for perturbed conditions different from training, and show that this framework unveils computational principles of how such neural dynamics enable flexible control of movement.

## 1 Introduction

Behavior arises from interactions among functionally and anatomically distinct entities of the brain, body, and physical environment. During voluntary movements, the central nervous system (CNS) transforms sensory and internally generated feedback into the muscle excitations required for proper movement execution. However, the underlying computations and dynamical strategies executed by the motor cortex (MC) during this transformation remain elusive. Much of the controversy lies around the proper placement of MC in the sensorimotor transformation and the corresponding inputs that it may receive from the upstream brain regions for pattern generation. Currently, there are two dominant theories regarding how the MC may accomplish this feat. The first is the representational perspective, which suggests that MC does not actively participate in pattern generation underlying sensorimotor transformation, and its activity relates to abstract movement-related representations, such as muscle features [1–11]. Alternatively, the dynamical systems perspective suggests that MC actively participates in generating outgoing muscle commands and its activity reflects the features of a dynamical system required to execute appropriate sensorimotor transformations [12–16]. The dynamical systems perspective can account for inconsistencies in the representational perspective, such as neurons that do not appear to code for any relevant movement or task variables [17]. Additionally, this theory provides a new lens through which to interpret MC activity, with analyses of population dynamics and their dependence on continuous sensory feedback together with the inputs from interacting brain regions, playing a key role [16, 18, 19].

This new perspective on MC activity requires new computational models to further our understanding of the complex transformations required during movement generation. Recently, task-driven or goal-driven models have risen in popularity due to their success in faithfully representing the high-dimensional population dynamics that generate behavior. These models are trained to perform a motor task considered analogous to that performed by a biological population of neurons, and their resulting solution is compared to neural recordings after training [20–22]. However, in these formulations, MC is usually considered a quasi-autonomous dynamical system that transforms condition-specific inputs into the corresponding muscle commands. The models in [20, 21] are thus trained to transform condition-specific inputs into the corresponding representations of muscle commands. In [22], a modular RNN-based MC model is trained to transform visual features of the sensory feedback into the inferred muscle velocities required to execute the movement. Despite their success, these models lack important components of the sensorimotor transformation: 1. They are not trained on real-time continuous sensory feedback, especially proprioception, 2. They do not interact with complex musculoskeletal dynamics, 3. They are not subject to physical laws of the environment, and 4. They are often trained on a very small subset of features that MC may consider as input or output. Therefore, these models fail to disentangle the role of sensory feedback from condition-specific inputs to MC. Importantly, these models fail to capture important features of cortical activity, such as the orthogonal subspaces observed during preparatory and movement periods for delayed reaching tasks [23]. Moreover, these models do not generalize to unseen movements. As a result, these models cannot be used for the prediction of cortical activity during these unseen conditions. For example, once a movement is learned, we have the ability to produce this movement arbitrarily fast or slow, within a range, while maintaining accuracy. Building on previous studies [20–22, 24], we show that building in the biological realism of the musculoskeletal dynamics for accurate sensory feedback generation (especially proprioception) and transforming the loss function in kinematics space towards goal-driven models of MC can vastly improve generalization and help better capture neural dynamics underlying movement generation.

Here, we articulate the point of view that motor pathways in the brain act as a feedback controller for the musculoskeletal system and drive optimal behavior under appropriate constraints [25–34]. Importantly, optimal feedback control (OFC) theory has elegantly highlighted the dependence of the CNS on feedback processes for movement generation under appropriate behavioral-level constraints [35–47]. These feedback processes include external sensory feedback (especially proprioception) and internally generated feedback through efference copies. Moreover, these studies suggest that cortical activity is optimized for musculoskeletal dynamics [16, 48]. As highlighted by OFC theory, since CNS transforms sensory feedback during stereotypical behaviors into muscle excitations, the resulting neural dynamics may also be predictable. However, previous models lack flexible neural network implementations of the controller, and fail to generate single-neuron predictions. Moreover, neural constraints that arise from the evolutionary standpoint cannot be implemented using these models, such as minimization of neural firing rates. Such neural constraints may be important for the explanation of neural dynamics and behavior as shown in [20].

Recently, deep reinforcement learning (DRL) has emerged as a promising way to train controllers for musculoskeletal models in highly complex tasks [49–55]. With DRL, it is possible to train these models using high-dimensional sensory feedback to produce muscle activations that generate complex movements, such as quick turning and walk-to-stand transitions [56]. The resulting behavioral-level dynamics have been shown to resemble experimentally recorded data [56]. Additionally, a recent approach called MotorNet [57] includes training controllers with differentiable biomechanical models; however, the correspondence of the resulting controllers to the MC or generalization to unseen conditions has not yet been well established.

We propose a computational framework, here termed muSim, to generate behavioral-as well as population- and single-unit-level neural predictions for unobserved movements and demonstrate the neurophysiological plausibility of our trained controllers. We introduce biological realism, such as the generation of accurate proprioception, into our computational framework through a sensorimotor loop implementation, incorporating anatomically accurate musculoskeletal models. This feedback loop is simulated using well-maintained physics simulation engines, further adding to their physical realism. Our modular framework allows the user to input musculoskeletal models of different animal species along with kinematics and other optional experimental data. The choice of training methodology, such as DRL algorithms, is also available. We provide various post-training analyses modules to analyze the population- and single-unit-level features of the trained network activity and to provide its qualitative and quantitative comparisons with neural data. Lastly, we implement different perturbation modules, including sensory feedback and neural perturbations, that enable robust hypothesis testing and generation, and prediction of dynamical mechanisms shaping the population-level activity patterns of MC.

Next, we validate muSim’s ability to capture the principle dynamical features apparent in MC activity for seen and unseen behavioral conditions. To do so, we train recurrent neural networks (RNNs) to control the complex musculoskeletal models in a goal-driven manner using DRL. We directly replicate the kinematics of macaque and mouse performing cycling and reaching movements with their forelimbs, and subsequently utilize our provided post-training analyses modules for comparisons with the corresponding neural data. We discover that for both seen and held-out behavioral conditions, the RNN solution faithfully aligns to the MC recordings at both the population- and single-unit-level. Compared to baseline models, such as open-loop RNNs and models built on movement parameters such as muscle activity and kinematic trajectories, our feedback-constrained model more closely resembles MC data across tasks and species, especially for unseen conditions. muSim’s ability to faithfully align to neural data across species, tasks, and behavioral conditions demonstrates the vital role that flexible sensory feedback transformations play during movement generation, and allows us to further probe the underlying principles of this computation using perturbation experiments.

## 2 Results

### 2.1 Development of goal-driven sensorimotor framework with musculoskeletal dynamics

Experimental and modeling studies have identified that motor cortical dynamics can be captured by a quasi-autonomous dynamical system, suggesting that its interaction with the rest of the neural circuit is not necessary for movement generation [12, 13, 20]. However, recent modeling and neurophysiological advancements investigating cortical connections with interacting neural circuits, such as sensory processing regions (especially proprioception), suggest that motor cortical dynamics are strongly modulated by feedback mechanisms [19, 58–60].

To obtain a model of MC based on feedback mechanisms, we develop a goal-driven computational framework, muSim. muSim is based on the modular implementation of the sensorimotor loop, i.e., transformation of sensory signals into muscle excitations and vice versa (Fig. 1a). muSim consists of various independent, interactive and interchangeable modules (See github page https://github.com/saxenalabneuro/muSim). muSim mimics the biological sensorimotor control loop where the controller, representing the MC, receives real-time sensory feedback and delivers muscle excitations to drive the musculoskeletal model producing diverse movements in the physics simulation environment (Fig. 1a). The sensory feedback can consist of proprioceptive feedback (muscle lengths/velocities and muscle forces), visual feedback (limb and target kinematics), joint positions/velocities and/or experimental stimulus data. The control inputs to the musculoskeletal model can consist of either muscle excitation signals or joint torques. Additionally, the introduction of Gaussian noise and time delays in the sensory feedback and control inputs can also be specified.

**Fig. 1.**
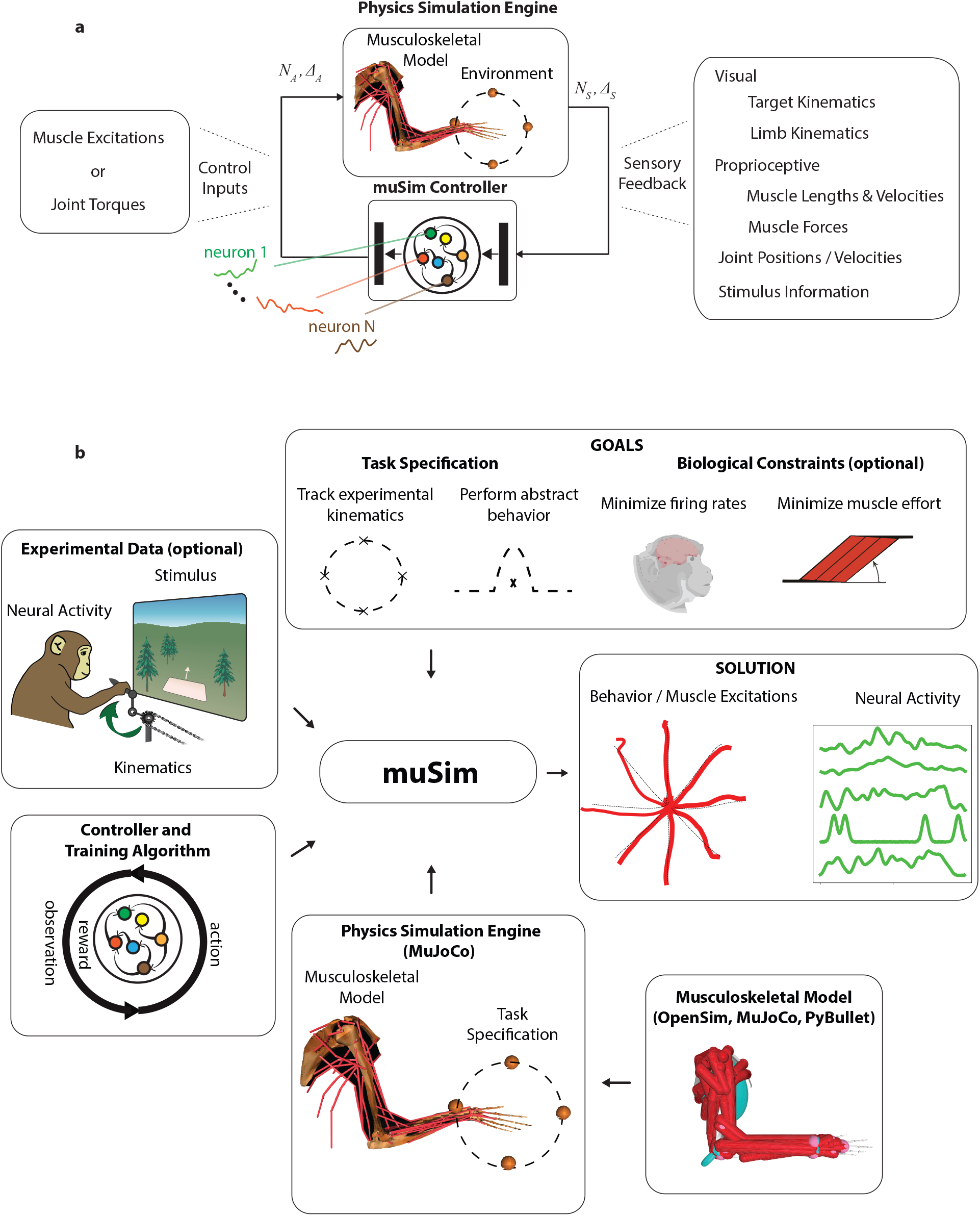
Development and overview of the goal-driven sensorimotor framework. **a**. In muSim, we build a biologically accurate model of the sensorimotor feedback loop. It consists of an anatomically accurate musculoskeletal model that can interact with its environment in a physics simulation engine. The physics simulation engine receives control inputs (muscle excitations or joint torques), executes the movement and generates the resulting sensory feedback. To execute a desired movement, the transformation from sensory feedback to control inputs is represented using a neural network controller. Here, the muSim controller consists of an RNN layer representing the model of MC and two feedforward layers representing the processing upstream and downstream of MC. Control inputs noise (*N*_*A*_) and delays (Δ_*A*_) as well as sensory noise (*N*_*S*_) and delays (Δ_*S*_) can also be specified. **b**. The proposed goal-driven sensorimotor framework consists of various independent interactive input modules: musculoskeletal model, physics-simulation engine, controller and training algorithm, experimental data, and goals specification. The anatomically accurate joint/muscle -based musculoskeletal model can be defined in widely used and well-maintained softwares, such as OpenSim, MuJoCo or PyBullet. The defined musculoskeletal model may require conversion for use in inter-compatible physics-based simulation engines, such as MuJoCo. The physics simulation engine additionally requires environmental and task specifications, such as experimental kinematics or conditions. The training algorithm is used for training the weights of the muSim controller to transform sensory feedback into control inputs that achieve the specified goals. The experimental data, such as kinematics, recorded neural-activity or stimulus data, can be used for task-specification to train the controller to reproduce recorded movements under physical experimental conditions and constraints. The goal specification consists of two sub-modules: task specification and optional biological constraints. The task specification can be based on experimental conditions, such as track recorded kinematics, or abstract behavior, such as obstacle avoidance. Additionally, biological constraints, such as reproduction of recorded neural activity on observed conditions or minimization of neural firing rates, can also be enforced. As output, muSim then produces the control inputs and the controller’s activity required to achieve specified goals, such as experimental kinematics, under the specified biological constraints.

muSim consists of several modules: an independent and interactive *controller, training algorithm*, physics simulation engine-based *environment, musculoskeletal model, experimental data*, and *goals-specification* (Fig. 1b). Modularity allows flexible integration of user-defined musculoskeletal models and training algorithms with minimal modifications. muSim can thus be easily adapted for diverse use-cases, such as simulating realistic environmental conditions, building neural and behavioral constraints, and using musculoskeletal models from different animal species.

Experimental data, such as recorded kinematics for different body parts or markers, single unit firing rates, and stimulus data, can be used to guide the training algorithm to achieve various goals. The goals-specification module utilizes the provided experimental data for task specification, such as tracking experimental kinematics. While training the model, constraints such as minimizing the neural firing rates can be specified to incentivize convergence to more biologically plausible solutions. These tasks are then simulated in the physics simulation engine, such as MuJoCo or PyBullet, with the musculoskeletal model receiving control inputs from the muSim controller. The specified training algorithm then trains the muSim controller to transform the continuous sensory feedback into control inputs that drive the musculoskeletal model to achieve the specified goals.

Here, we demonstrate different aspects of muSim using a validated macaque limb model with 39 muscles [61], which was adapted from OpenSim to MuJoCo as in [62, 63] for computational efficiency (see Methods). We additionally test our framework on a mouse musculoskeletal model with 18 muscles, implemented in the PyBullet physics simulation engine (see Methods). Our task specification utilizes recorded experimental kinematics for tracking different movements, such as reaching and cycling in macaques and alternation in mice. We also simulate delayed reaching to analyze the controller’s preparatory responses. We use an RNN architecture for the controller representing the recurrent connections of the biological MC. The controller consists of three layers, an RNN layer representing the model of MC, along with two feedforward layers representing upstream and downstream interacting neural circuits.

To summarize, we have introduced a computational framework, muSim, to provide a comprehensive model of the sensorimotor loop. Our modular implementation enables flexible integration of user-defined experimental task specifications and musculoskeletal models to facilitate comparison of the trained controller’s activity with the recorded neural data. We also provide modules to integrate user-defined training algorithms and feedback specifications, in addition to the provided DRL training algorithms.

### 2.2 muSim controller achieves high kinematic accuracy and generalizes across conditions in a variety of motor tasks

We use DRL to train the muSim controller to produce either the user-specified experimentally recorded kinematics, or the tasks themselves under specified constraints (see Methods) (Fig. 2a). We provide inverse kinematics and evolutionary algorithms to figure out the initial pose of the musculoskeletal model as specified by the user and to determine if the user-specified movement is feasible (see Methods). DRL then trains the muSim controller to maximize a given reward function under specified neural- and kinematics-level constraints. Here, we adapted the soft actor-critic (SAC) algorithm with added neural regularizations to train the muSim controller. These neural regularizations also enabled the muSim controller to capture biological interpretability in the neural and kinematics subspaces, such as cortical dynamical strategies and generalization to unseen conditions different from training. The critic networks associated with reward function learning may be considered a proxy to other areas of the brain’s reward system, such as the ventral striatum, and may reflect dopaminergic projections to MC [64].

**Fig. 2.**
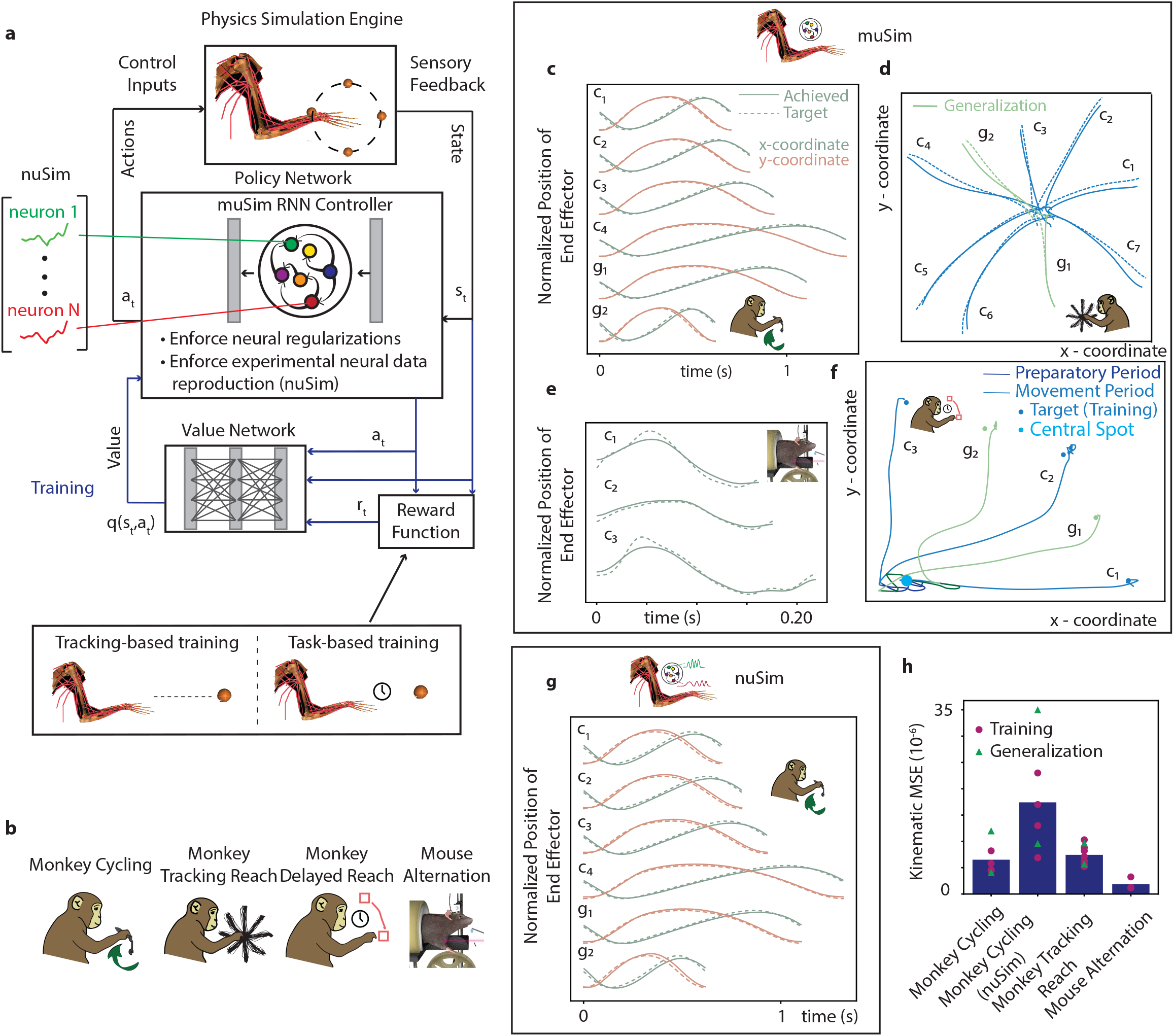
muSim achieves high kinematic accuracy on various tasks trained using DRL. **a**. While training, various neural regularizations, such as minimization of neural firing rates, can be enforced on the muSim controller/policy network. Additionally, in nuSim, a subset of the muSim controller’s single units can be constrained to reproduce the firing rates of experimentally recorded neurons during the task. The value network outputs the q-value of state and action pair that represents its cumulative reward. In tracking-based training, the reward is generated at each timestep as the target position varies from initial to final position usually following the experimentally recorded kinematics. In task-based training, experimental kinematics may not be required and the task-specification can be relatively more abstract, such as delayed reach, making the reward relatively sparse. Blue connections are used only during the training, while the others (black) are used during both the training and testing. **b**. We train muSim for various tasks: monkey cycling, monkey tracking reach, monkey delayed reach and mouse alternation tasks. We use anatomically accurate musculoskeletal models for the macaque and the mouse (see Methods). **c**. muSim achieves high kinematic accuracy on training conditions, (*c*_1_ - *c*_4_). muSim generalizes to testing conditions unseen during training, *g*_1_ and *g*_2_. Interpolated condition (*g*_1_) lies in the distribution of the training conditions, while the extrapolated condition (*g*_2_) lies out-of-distribution of training conditions. **d**. muSim achieves high kinematic accuracy for the monkey tracking reach task for the seven relatively straight training conditions while generalizing to two unseen testing conditions. **e**. muSim achieves high kinematic accuracy for the mouse alternation task for the three training conditions. **f**. muSim achieves high kinematic accuracy for the monkey delayed reach task in terms of fixating at the central point during the preparatory period and performing the actual reach during the movement period. **g**. Same as **c** but for nuSim. **h**. The MSE between the target and achieved kinematics for various tasks is plotted for all the seen training (circle) and unseen generalization (triangle) conditions. The bars show the mean kinematic MSE across all the training and generalization conditions for the given task (x-axis).

In this formulation, DRL consists of a policy and a value network. The policy network represents the muSim controller and transforms the sensory feedback into the required muscle excitations, here representing the motor pathways. During training, the policy network’s weights are updated to maximize the cumulative reward as specified by the reward function *r*(*s*_*t*_, *a*_*t*_), while optionally being constrained to follow neural constraints and regularizations (see Methods). In addition to task specification, reward function can be used to enforce biological constraints, such as minimization of muscle effort.

Two recent approaches have shown promise in obtaining predictive models of neural circuits: data-driven modeling and goal-driven modeling [65]. In data-driven modeling, biophysically-detailed or statistical models are fit directly to the recorded neural data. The trained models are then probed for neural predictions and insights into the developed circuit properties [66, 67]. In goal-driven modeling, neural data is not directly used for training the model; instead, the model is trained on a task considered analogous to that performed by the neural circuits. The trained model’s activity is then compared with the recorded neural data to generate predictive insights [20, 21, 68]. The implementation of the sensorimotor loop in muSim enables simultaneous goal- and data-driven modeling to further constrain the solution space of the trained controller to develop a direct correspondence with the recorded single units. The resulting constrained muSim is denoted as nuSim. We develop a dynamical systems model of MC that is both goal- and data-driven: it transforms sensory feedback into muscle excitations that are converted to kinematics, while directly encoding recorded MC single units. During training, we constrain a subset of the nuSim RNN’s nodes to the empirically recorded MC single unit activity only for the training conditions (see Methods).

The reward function can be used to specify the training using DRL as either tracking-based or task-based training. For tracking-based training, the muSim controller is trained to reproduce experimentally recorded limb movements. An immediate reward *r*(*s*_*t*_, *a*_*t*_) is generated based on how well the musculoskeletal model’s limbs follow the experimentally specified movement at each timestep *t*. On the other hand, task-based training can be used to model the task itself without requiring the experimental kinematics at each timestep. Here, the reward function *r*(*s*_*t*_, *a*_*t*_) tends to be sparse. In such cases, curriculum learning can be used (see Methods). In curriculum learning, the controller is gradually trained to build up a rich repertoire of movements of increasing difficulty.

Using our computational framework, muSim, we trained the controller for four different tasks (Fig. 2b). It is important to note that the recorded neural data was not used at all during muSim training.

1. Monkey Cycling Task: An adult male rhesus macaque was trained to produce a cyclic movement of the forelimb at 8 different speeds, while kinematics and single-unit activity from premotor and motor cortex were recorded [21]. We used experimentally recorded kinematics during a subset of four different speeds as a task specification for muSim to train the controller, withholding other speed conditions for testing purposes. We used an anatomically accurate musculoskeletal model of macaque (see Methods) [61, 69].
2. Monkey Tracking Reach Task: We trained the muSim controller on a center-out reaching task with relatively straight reach trajectories to the outer targets (see Methods) [13].
3. Monkey Delayed Reach Task: The delayed reach task was designed to consist of preparatory and movement periods. During the preparatory period, the muSim RNN has to fixate the musculoskeletal model’s limb at a central fixation point while preparing the reach to the displayed outer-target. A go-cue then marks the transition to the movement period, and the muSim RNN executes the reach to the outer target. We first use curriculum learning to stabilize the DRL training for the delayed reach task. In curriculum learning, the muSim RNN is first trained to perform reaches to the outer targets from a central point using tracking-based training (without a preparatory period). Subsequently, the preparatory period is introduced and the task-based training ensues. This curriculum can be flexibly adjusted to account for experimental strategies for training macaques on tasks (see Methods).
4. Mouse Alternation Task: We trained an anatomically accurate mouse musculoskeletal model (see Methods) [70] to reproduce an alternation behavior (akin to cycling) performed by mouse forelimb, as in [71] (see Methods).

Finally, the nuSim controller was trained on the monkey cycling task with a subset of the policy network’s units constrained to the experimentally recorded single-unit firing rates. The trained muSim and nuSim controllers achieved high kinematic accuracy for all training tasks with relatively low mean squared error (MSE) between the experimental and simulated trajectories (Figs. 2c - 2g). The kinematic MSE is quantified in Fig. 2h. The trained muSim/nuSim also generalized to conditions unseen during training for various tasks in the kinematics space (Fig. 2c - 2g). For the monkey cycling task (Figs. 2c and 2g), the unseen interpolated condition *g*_1_ lies in the distribution of the training speeds, whereas the unseen extrapolated speed condition *g*_2_ lies outside the distribution of the training speeds.

### 2.3 muSim RNN dynamics mimic recorded neural population dynamics for sensorimotor tasks

After validating that the trained muSim/nuSim controllers achieve high training and generalization kinematic accuracies, we first qualitatively compare the resulting controller’s activity with the recorded neural data. Given the intrinsic (for example, connectivity) and extrinsic (for example, motor task) constraints to the neural circuits, the most informative and meaningful states of the neural data occupy a low-dimensional manifold spanned by a few states far lower than the recorded single-units. This gives rise to the general idea of population-level dynamics. Dimensionality reduction methods, such as principal component analysis (PCA), jPCA and fixed-point structure, have commonly been used to identify population-level dynamics and strategies for the control of movement. Here, we use these methods to assess the dynamical alignment between the population-level dynamics underlying the controller’s activity and the neural data for the trained motor tasks (Fig. 3a).

**Fig. 3.**
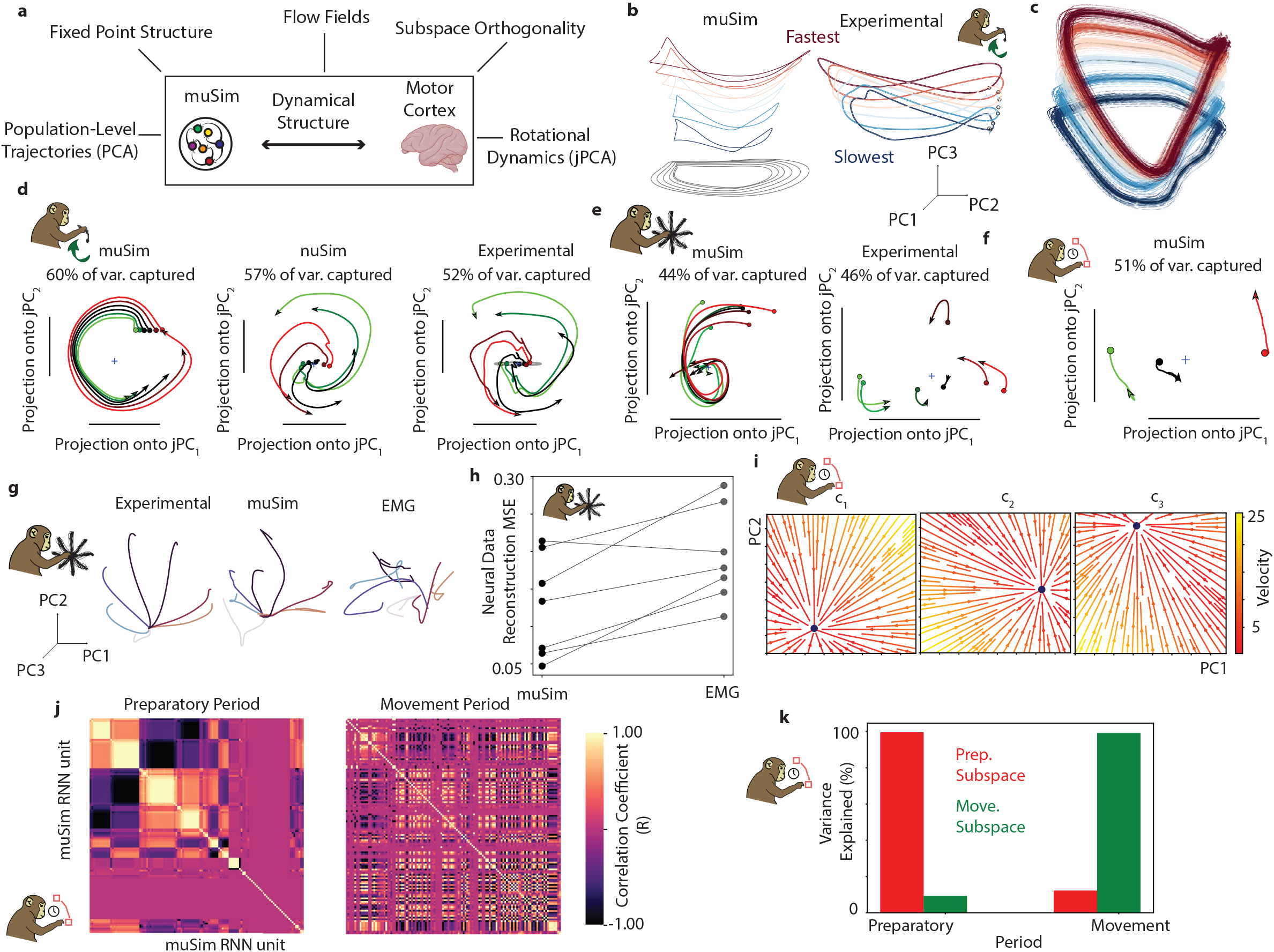
Network dynamics mimic the recorded neural dynamics for sensorimotor tasks. **a**. Various post-training analyses modules based on dimensionality reduction methods, such as PCA, jPCA and fixed point analysis, are provided to qualitatively assess the dynamical alignment of muSim RNN with the empirically recorded neural data. **b**. (Left Panel) muSim RNN population-level trajectories in the top 3 PCs subspace for the training and generalization (testing) conditions for the monkey cycling task. The trajectories for various speeds show a stacked elliptical structure. (Right Panel) Corresponding experimental neural trajectories in the top 3 PCs subspace. **c**. muSim RNN trajectories exhibit stable limit cycles. The sensory feedback to the muSim controller was perturbed to start from a different state. The black lines represent the original unperturbed muSim RNN trajectories corresponding to different speed conditions. Each arrow points to the direction muSim RNN controller activity converged to after perturbation. All the perturbed trajectories converged towards the original limit cycle corresponding to each speed condition. **d**. jPCA trajectories for different speed conditions (different colors) for the monkey cycling task. jPCA shows the presence of strong rotational dynamics for muSim (left panel), nuSim (middle panel) mimicking the experimental neural data trajectories (right panel). Each trace shows the evolution of the state trajectory over 620 ms after movement onset. **e** and **f**. Same as **d** but for the monkey tracking reach and delayed reach tasks during the movement period, respectively. **g**. Structure of experimental neural trajectories (left panel) for the seven reach conditions (different colors) in the top 3 PCs subspace for the monkey tracking reach task after controlling for the delay period. LRA was used to test the generalization of muSim and EMG trajectories for each speed condition. After LRA, the resulting trajectories are shown for muSim (middle panel) and EMG (right panel). muSim trajectories for different speed conditions followed a similar pattern as the experimental trajectories. EMG trajectories did not show any organized pattern. **h**. For the monkey tracking reach task, MSE between the muSim and experimental trajectories and between EMG and experimental trajectories after LRA in top 8 dimensional PCs subspace is shown. **i**. Flow field analysis in the top 2 PCs subspace shows that the muSim RNN controller exhibits condition specific fixed-point (attractor) structure during the preparatory period. For the flow field analysis, the sensory feedback to muSim RNN controller in the original high-dimensional subspace was perturbed to start from different state during the preparatory period. The resulting muSim RNN activity converged to the condition-specific attractor after perturbation. **j**. Preparatory period (left) and movement period (right) correlation matrices for the muSim RNN units. Each entry in the matrix represents the degree to which the response was similar for the two units during that period. **k**. Percentage of variance captured by the preparatory (red) and movement (green) subspaces during the two epochs. The muSim RNN response during each epoch was projected onto the top ten dimensional preparatory and movement subspaces. The left pair of bars corresponds to the variance captured during the preparatory period. The right pair of bars corresponds to the variance captured during the movement period.

We use a linear dimensionality reduction method, PCA, to identify the population-level dynamics as dominant covariation patterns across the single-units of muSim RNN collectively for different conditions of the trained motor task [72]. The resulting population-level trajectories spanned by the dominant covariation modes can be visualized to identify the population-level structure across the training and generalization conditions of the motor task. For the monkey cycling task, the MC population structure exhibits stacked elliptical neural trajectories (Fig. 3b, right panel) having low trajectory tangling and separated along a speed dimension to enable flexible control of speed [21]. We found that muSim controller employs similar population-level dynamical strategies for this task with stacked elliptical trajectories separated mainly along a speed dimension (Fig. 3b, left panel).

Next, to find out whether the muSim population-level trajectories for different speed conditions exhibit stable limit cycles, we perturbed the muSim RNN trajectories (in the high-dimensional single-unit state space) to start from a different state at each timepoint during the cycle. The perturbed trajectories recovered towards the original unperturbed state at each timepoint, further suggesting a stable limit cycle for each speed condition (Fig. 3c). This finding also points to the existence of a well-tuned feedback control policy underlying muSim controller that contributes to its robustness to perturbations.

However, one limitation of PCA is its inability to clearly capture the rotational dynamics in the resulting population-level trajectory spanned by dominant neural modes. We used jPCA to quantify such rotational dynamics in the data [13]. This tool also quantifies the distribution of angles between the state and its derivative. Pure oscillatory dynamics result in angles distribution close to 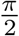 . MC population activity has been shown to exhibit rotational dynamics for various rhythmic and non-rhythmic movements [13, 20]. Such quasi-oscillatory patterns may reflect the underlying movement generating dynamics present in the population response that may not represent the movement parameters. For the monkey cycling task, jPCA revealed the presence of strong rotational dynamics in muSim population activity for all speed conditions, with rotational dynamics explaining a significant variance of the RNN activity for both muSim and nuSim, mimicking the experimentally observed rotational dynamics (Fig. 3d). The mean of distribution of angles between the muSim state and its derivative lies close to 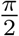, indicating the presence of a strong rotational component. Similarly, for the monkey reaching and delayed reaching tasks, jPCA revealed the presence of strong rotational dynamics for all reach conditions during the movement period (Figs. 3e and 3f, respectively).

Associative studies have examined the rich fixed point structure in network computations to elucidate the dynamical strategies underlying the network’s population-level response and link them to the relationship between task conditions [73]. Following a similar approach, we computed the stable (attractor) and unstable (repeller) fixed points implemented by the muSim RNN after solving the sensorimotor task. Such fixed point structure has yielded insightful interpretations of dynamical strategies employed by RNNs driven mainly by autonomous dynamics, especially in cognitive tasks [74]. However, for the monkey cycling task, the fixed point structure contained a single condition-independent stable fixed point (Supplementary Fig. 1), and was not informative of the dynamical strategies underlying the muSim RNN population response. For this particular task, the lack of a rich fixed-point structure suggests the reliance of muSim RNN on sensory feedback as opposed to being driven mainly by internal recurrent (autonomous) dynamics. This is also validated by our results on the selective feedback ablation below indicating that sensory feedback ablation has a more drastic effect on the muSim kinematic accuracy as compared to the recurrent connections ablation (see Section 2.6).

The dimensionality reduction methods discussed above do not consider the time-dependent relationship between the muSim RNN and neural population-level trajectories when computing the dynamical alignment. We used linear regression analysis (LRA) to compute the population-level trajectories for the muSim RNN while taking into account their time-dependent relationship with neural trajectories (see Methods). For the monkey tracking reach task, we then used LRA to compare the neural trajectories against the muSim RNN and EMG trajectories for the different reach conditions in the top 8 dimensional PC subspace. After LRA, muSim-RNN population trajectories more closely resembled the neural population trajectories, whereas EMG trajectories failed to capture the population-level structure present in the neural data (Fig. 3g). muSim trajectories also exhibited lower MSE with the experimental neural trajectories compared to the EMG trajectories (Fig. 3h). These findings collectively suggest that population-level MC and muSim RNN trajectories primarily contain dynamical features required for movement generation and may not represent movement features, such as muscle activity.

Next, we examine the muSim RNN preparatory activity for the delayed-reach task. The optimal subspace hypothesis postulates that MC population activity during the preparatory period travels to an optimal region of state space from which to generate the given reach [75]. The initial condition hypothesis further refines the optimal region and suggests that the preparatory activity sets the initial condition for the movement generating dynamical system [12]. We leveraged the robustness of muSim to perturbations to investigate if a specific initial condition is attained during the preparatory period. For this, we analyzed the fixed-point (attractor) structure for different reach conditions during the preparatory period for the monkey delayed-reach task. We perturbed the muSim-RNN trajectories in the high-dimensional single-unit state-space during the preparatory period for the flow-field analysis in Fig. 3i. Given sufficient preparatory time, all muSim trajectories converged back to the original condition-specific fixed-points after perturbation. Therefore, muSim exhibited a strong condition-specific fixed-point (attractor) structure. This finding suggests that a condition-specific initial state for the movement generating dynamical system is reached. During the movement period, the trajectories then showed rotational dynamics evolving from these fixed-points (Fig. 3f).

As shown above, the muSim RNN’s population response during the preparatory period mainly develops to set the initial state for the subsequent movement period dynamics. During the movement period, the underlying population response reflects drastically distinct computations governed by rotational dynamics required for movement generation. How do the population-level signatures of the same set of neurons (muSim RNN units) that subserve two distinct computations change across preparatory and movement periods on a very short timescale? To address this, we first calculated the correlation matrices for the muSim-RNN population for the preparatory and movement periods. To aid visualization, we kept only those units that showed significant variance (*>* 0.3) either during the preparatory period or the movement period. The subsequent analysis and results remained similar even if all units were included. The ordering of units was chosen to highlight the structure (using hierarchical-clustering) during the preparatory period and was kept the same for the movement period (Fig. 3j). The correlation structure changed markedly across the preparatory and movement periods, consistent with the experimental studies [23].

As indicated by the perturbation studies, there exists a condition-specific attractor structure that governs the initiation and subsequent evolution of movement generation dynamics. This shows that there exists some lawful relationship between the population responses across the two epochs. However, the marked change in the correlation structure shows that the set of neurons with shared properties changes rapidly across the epochs. These findings suggest that population responses during the two epochs are linked but occupy two different and orthogonal subspaces to subserve the distinct computations during the two epochs. To test for the hypothesis that the preparatory epoch dimensions are orthogonal to the movement epoch dimensions, we used PCA to project the population response onto the two epochs separately. As done in [23], we performed PCA separately on the preparatory epoch responses to find the preparatory-epoch principal components (prep-PCs) and similarly on the movement epoch responses to find the movement-epoch principal components (move-PCs). Top 10 prep-PCs captured the highest variance of the preparatory epoch responses and the top 10 move-PCs captured the highest variance of the movement epoch responses. However, prep-PCs captured very little movement epoch data variance and move-PCs captured very little preparatory epoch data variance (Fig. 3k). This finding indicates that preparatory epoch responses are orthogonal to the movement epoch responses. This is significant because the existing dynamical systems models of MC (based on quasi-autonomous dynamics) can not capture the orthogonality between the preparatory and movement-period population responses [20]. To further investigate the role of sensory feedback in orthogonality, we then used sensory feedback and EMG from the muSim controller trained for the monkey delayed-reach task to train an open-loop (OL) RNN. Interestingly, we did not find any structured pattern in the correlation matrices during the preparatory and movement epochs (Supplementary Fig. 2a). OL RNN also did not develop orthogonal subspaces across the two epochs (Supplementary Fig. 2b). This finding suggests that other aspects of muSim, such as neural regularizations and exploration, likely work in synergy with with sensory feedback features towards the emergence of orthogonal subspace structures. Lastly, to aid future population-level analyses such as PCA, and to test the dynamical alignment between trained muSim RNN and neural data, we have provided post-training analysis modules in the framework.

### 2.4 muSim enables generalizable decoding of recorded neural activity from the MC

Next, we used three different methods to directly compare the trained muSim and recorded neural data, and quantify the resulting similarity. These methods included canonical correlation analysis (CCA) for quantifying the similarity between population level responses (see Methods). These methods also included LRA and Procrustes for quantifying the similarity between single-unit level responses (Fig. 4a) (see Methods). First, we investigate the similarities for the monkey cycling task.

**Fig. 4.**
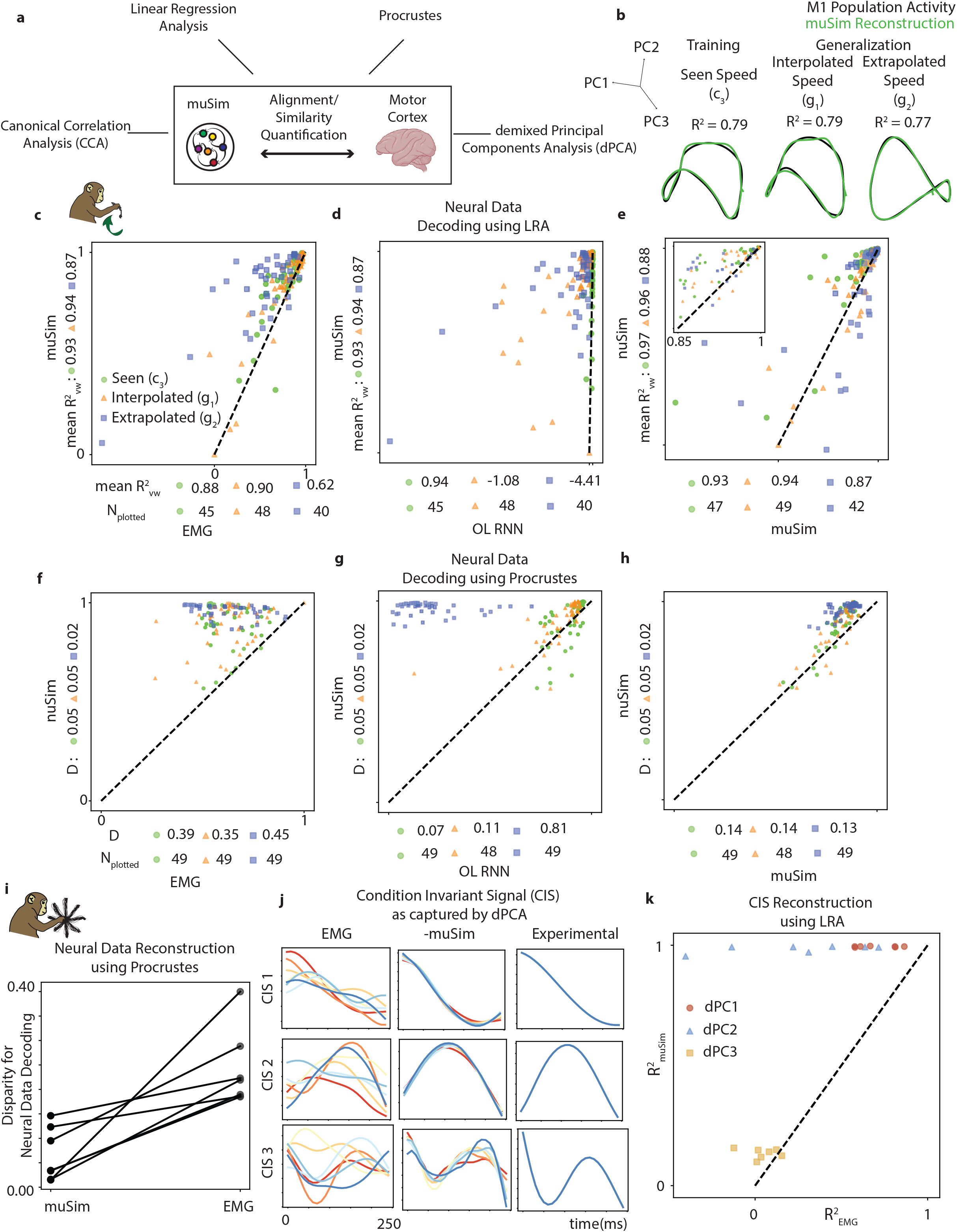
muSim outperforms the goal-driven and representational models of MC in terms of the decoding accuracy while achieving generalization. **a**. We provide various post-training modules, based on CCA, Procrustes, LRA and dPCA, to quantify the similarity between the muSim RNN trajectories and neural data. **b**. The first 3 PCs of MC population activity (black trajectories) reconstructed from the muSim RNN activity (green trajectories) using the inverse canonical correlation analysis (CCA) for the seen (left panel), interpolated (middle panel) and extrapolated (right panel) speed conditions. We observed a high mean *R*^2^ value between the actual and reconstructed MC population activity in the top 10 PCs subspace for all speed conditions. **c**. Scatter plot shows the comparison of LRA-based single neuron decoding accuracy, as given by variance-weighted mean 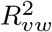, of muSim against the EMG-based representational model of MC on the seen, interpolated and extrapolated speeds. For each neuron (dot), the *R*^2^ of muSim RNN is shown along the vertical axis and *R*^2^ of the alternative model (EMG) is shown along the horizontal axis. If a neuron lies above the dashed unity line, it is better predicted by the muSim as compared to the alternative model. Only those neurons are shown for which the *R*^2^ ≥ 0 for either of the two models. The total number of neurons plotted *N*_*plotted*_ for each condition are also indicated.The variance-weighted mean 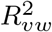 values between the experimentally recorded firing rates and their reconstructions are also shown. Inset shows the zoomed-in version of the outer scatterplot. **d** and **e**. Same as **c** but for comparison against the goal-driven OL-RNN and goal- and data-driven nuSim models. **f**. Scatter plot shows the comparison of procrustes-based single-neuron decoding accuracy (*R*^2^) of nuSim against the EMG-based representational model of MC on the seen, interpolated and extrapolated speed conditions. For each neuron (dot), the *R*^2^ of nuSim RNN is shown along the vertical axis and *R*^2^ of the alternative model is shown along the horizontal axis. Only those neurons are shown for which the *R*^2^ ≥ 0 for either of the two models. The total number of neurons plotted *N*_*plotted*_ for each condition are also indicated. The disparity (D) values between the experimentally recorded firing rates and their reconstructions are also shown. **g** and **h**. Same as **f** but for comparison against goal-driven OL-RNN and neural-unconstrained muSim. **i**. Disparity values for muSim and EMG for recorded neural data reconstruction for 7 reach conditions for monkey tracking reach task. **j**. For the monkey tracking reach task, top 3 condition invariant signals (CIS) as captured by the dPCA for the seven reach conditions (different colors) for the EMG, muSim RNN and experimentally recorded neural trajectories. **k**. For monkey tracking reach task, generalization performance of muSim RNN vs EMG for reconstructing CIS across various reach conditions and dPCs.

CCA has been extensively used to quantify the similarity between RNN models and neural data [20, 76, 77]. In short, CCA finds the weightings for individual units in the two datasets to find the underlying common patterns. The resulting two sets of maximally correlated reweighted data sets are known as canonical variables. The mean canonical correlation value (*R*^2^) that we report gives the mean correlation across the the first 10 canonical variables. CCA showed a high correlation (*R*^2^) of 0.79 for the conditions that muSim was trained on (Fig. 4b, left panel). We then tested the alignment between the muSim-RNN activity and the neural data on the unseen interpolated and extrapolated speed conditions. CCA showed high *R*^2^ values of 0.79 and 0.77 for the interpolated and extrapolated speed conditions, respectively (Fig. 4b, middle and right panels). We found that 10 canonical variables captured almost 100% of the neural data variance. muSim thus generalized to unseen conditions in both the kinematic and neural population subspaces.

Next, we used LRA to quantify the similarity between single-unit level responses. Briefly, we trained a linear regressor from the muSim RNN’s activity to the recorded MC firing rates using data from all task conditions except the held-out condition. We then tested the trained linear model on the held-out condition to transform the muSim RNN activity into the neural data subspace (see Methods). The *R*^2^ value between two data sets is reported in the neural data subspace. We compared the single neuron decoding accuracy (*R*^2^) of the muSim-RNN to the existing goal-driven and representational MC models using LRA. We considered the following commonly used goal-driven and representational models for comparison [1, 2, 20, 21]: (1) Empirically recorded EMG to test if the trained controller provides a comparison against the representational view that the cortical activity represents muscle commands; (2) Goal-driven open-loop RNN trained to transform a scalar speed signal to corresponding experimental recordings of muscle signals (electromyography; EMG) to provide a comparison against existing goal-driven models of MC; (3) Neurally constrained nuSim to provide a comparison against the simultaneous goal- and data-driven MC model; (4) Experimentally recorded kinematics to test if the MC model outperforms the representational view that signals mirroring the variables relevant to the task can capture cortical activity. The muSim/nuSim RNN outperformed all alternative models in decoding accuracy for the generalization speed conditions (Figs. 4c-4e and Supplementary Fig. 3). The higher performance of muSim as compared to the EMG and kinematics-based model of MC stresses the importance of additional single-unit-level features that may not be captured by the cortical models based on the representational view. Furthermore, MC is usually considered a quasi-autonomous dynamical system modeled as a pattern-generating RNN that transforms condition-dependent sparse input signals into the corresponding muscle commands. The higher performance of muSim against these pattern-generating RNNs, such as OL RNNs, highlights the importance of sensory feedback features and further stresses that MC single-unit level activity is optimized for musculoskeletal dynamics. This finding is consistent with OFC models of MC [16].

We further investigated and quantified the similarity between single-unit level responses of the muSim RNN and neural data using procrustes analysis (see Methods). Briefly, procrustes analysis linearly transforms (mainly, scales and rotates) the muSim RNN activity to align it optimally with the single-unit responses in neural data. We report the *R*^2^ value in the neural data subspace after transformation by procrustes analysis. Procrustes analysis is a well-established technique for quantifying similarity on single-unit level and is a more stringent test than CCA and LRA [22, 78, 79].

We compared the single-unit decoding accuracy of the trained nuSim controller against the existing data- and goal-driven models of the MC. The scatter plots (Figs. 4f - 4h) compare the *R*^2^ per neuron between its reconstruction produced by an alternative model (horizontal axis) against that produced by the nuSim (vertical axis) after transformation by procrustes analysis. We see that for all the seen and unseen speed conditions tested, most of the neurons lie above the dashed unity line, which means that they are being better predicted by the goal- and data-driven nuSim model as compared to the alternative models. We also observed the lowest disparity D (ℒ_2_ norm between the network reconstruction and recorded single-unit activity) for the nuSim as compared to the alternative models. Importantly, nuSim outperformed muSim in decoding accuracy especially for the generalization (interpolated and extrapolated) task conditions. This finding suggests that constraining a subset of nuSim units to neural data for training conditions further restricted its solution space as compared to muSim. The constrained solution space of nuSim generalized such that the relationship between the individual single-units and task conditions was better conserved than muSim.

Next, we investigated if muSim outperforms alternative models in neural data decoding for a variety of other tasks. We used procrustes analysis to compare the decoding accuracy of muSim RNN against EMG-based model for the monkey tracking reach task. For all seven reach conditions, muSim showed relatively lower disparity *D* (see Methods) indicating better neural data reconstruction (Fig. 4i). Moreover, MC activity has been shown to exhibit a strong condition invariant signal (CIS) across different reaches that can be captured using the demixed PCA (dPCA) [80]. CIS has been shown to reflect the movement timing for the upcoming reach. For the monkey delayed reach task, CIS for different reach conditions as captured by dPCA for EMG, muSim and experimental data are shown in Fig. 4j. To quantify how well the muSim activity can capture the CIS, we used LRA to fit the CIS for the muSim activity with that of the recorded data for six of the seven reach conditions and tested the generalization performance (*R*^2^ between the muSim and recorded data CIS) on the held-out condition. To provide a comparison, we performed similar LRA analysis for the empirically recorded EMG signals. Fig. 4k shows that generalization *R*^2^ values for most of the reaching conditions lie above the dashed unity line, indicating that muSim activity has a well-defined CIS that is not present in the muscle signals. This finding suggests that muSim RNN can capture abstract dynamical features of neural data that may not be reflected in the movement parameters. Lastly, to aid future analyses seeking quantification of similarity between the trained muSim/nuSim RNN and neural data, we have provided *quantitative* post-training analysis modules.

### 2.5 Control studies show that kinematic accuracy and muscle dynamics are crucial to the decoding performance of muSim

Here, we develop control studies for the methods used to quantify the similarity between the muSim RNN and neural data. The results in this section are mainly reported for the monkey cycling task. The muscle-based muSim (see Methods) consists of anatomically accurate musculoskeletal model with joints actuated by muscles in the sensorimotor transformation. The joint-based muSim (see Methods) consists of a skeletal model with joints instead actuated by motor actuators. For the random-muSim, we used a randomly initialized controller for muscle-based muSim. First, we used PCA to compare the population-level trajectories for the muscle-based muSim against those of joint-based muSim and random-muSim. Importantly, population-level trajectories for the other models showed marked differences from the neural data. For the joint-based-muSim, the population trajectories become highly overlapped indicating an increase in the across-speeds trajectory tangling. Moreover, across-speeds separation between the population trajectories along the speed-axis decreased significantly (Fig. 5a). For the random-muSim, the population trajectories settled at condition(speed)-specific stationary points after the first few cycles (Supplementary Fig. 4a). We also used jPCA to quantify the variance captured by rotational dynamics in the population response (Supplementary Figs. 4b and 4c). For the joint-based-muSim, we found that the rotational dynamics captured slightly less variance than the muscle-based muSim. Moreover, the distribution of angles in jPCs plane showed bimodal distribution with increased contribution from angles greater than 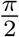. For the random-muSim, the population-level trajectories showed no rotational dynamics. Next, we used LRA to compare the *R*^2^ values between the network reconstruction and the neural data. Muscle-based-muSim outperformed other models, especially random-muSim, in terms of the reconstruction *R*^2^ values for the seen, interpolated and extrapolated speed conditions (Fig. 5b). Similarly, for the procrustes analysis, the muscle-based-muSim showed lower disparity values than other models for the seen, interpolated, extrapolated and concatenated speed conditions (Fig. 5c). The above results signify that muscle dynamics may play a significant role in the decoding accuracy of muSim.

**Fig. 5.**
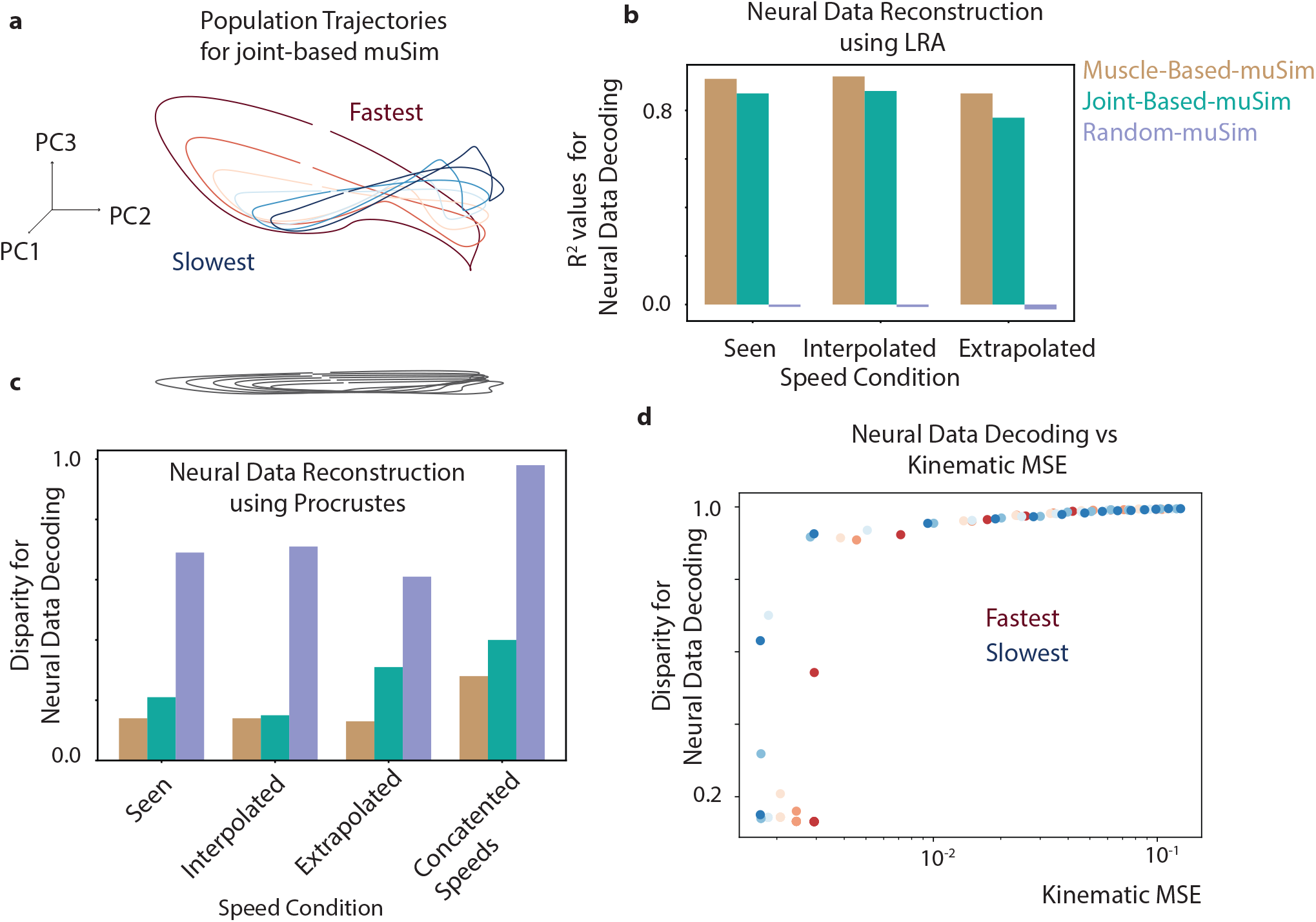
Control studies show that muscle dynamics and kinematic accuracy are crucial to the neural decoding performance of muSim. **a**. Joint-based-muSim RNN controller trajectories for the different speed conditions (different colors) in the top 3 PCs subspace for the monkey cycling task. The trajectories become distorted, as they are no longer elliptical and lose the separation along the speed axis, as compared to the experimentally observed trajectories. **b**. Bar plot comparing the LRA-based decoding *R*^2^ values for the muscle-based, joint-based and randomly initialized muSim for the seen (left bars), interpolated (middle bars) and extrapolated (right bars) speed conditions. **c**. Bar plot comparing the Procrustes-based decoding disparity values for the muscle-based, joint-based and randomly initialized muSim for the seen, interpolated, extrapolated and concatenated speed conditions. Concatenated speed condition represents the condition where the network and neural responses are separately concatenated in the time axis for all the seen and testing speed conditions. **d**. Scatter plot comparing the Procrustes-based disparity values against the kinematic MSE for the seen and unseen speed conditions. Lower kinematic MSE results in exponentially lower disparity values suggesting a higher alignment between the muSim RNN and experimentally recorded neural states for all speed conditions.

Does the kinematic accuracy of muSim determine its corresponding neural decoding accuracy? To test this, we replaced the trained muSim controller with a randomly initialized controller in 100ms time intervals and examined the resulting neural decoding accuracy. For CCA, the heatmap in Supplementary Fig. 6a shows a marked decrease in the *R*^2^ values if the trained-RNN is transitioned to random-RNN before the cycle for the corresponding speed condition is completed. For the procrustes analysis, Supplementary Fig. 6b shows that there is a marked increase in the disparity values if the trained-RNN is transitioned to random-RNN before the cycle for the corresponding speed condition is completed. This indicates that kinematic accuracy plays an important role when testing the similarity between muSim RNN and neural data using methods, such as CCA and procrustes analysis.

To further quantify the relationship between the muSim kinematic accuracy and its neural decoding accuracy, we examined the disparity values (*D*) for different speed conditions against the kinematic MSE for procrustes analysis. We observed exponentially increasing disparity values with the increasing kinematic MSE (Fig. 5d). Similarly, Supplementary Fig. 6c shows exponentially decreasing *R*^2^ values for different speed conditions with the increasing kinematic MSE value. This finding suggests that kinematic accuracy of muSim RNN plays an important role in its neural decoding accuracy.

### 2.6 Perturbation experiments using nuSim enables robust hypothesis testing and generation

Skilled mammalian behavior relies on neural circuits that generate robust and flexible patterns of population-level activity with complex underlying dynamical mechanisms. Decades of studies have linked specific patterns of population-level activity in MC to the generated behavior [13, 21]. However, association alone is not enough to uncover the exact nature of the dynamical mechanisms and computational principles governing the evolution of population-level patterns. Importantly, specific computational hypotheses will be regarded as falsifiable predictions if they cannot predict the computational and dynamical mechanisms (mainly, flow fields) shaping population-level activity. Perturbation experiments provide a criterion with which to evaluate such hypotheses more robustly than the mere association of population patterns with generated behavior. Here, we perform perturbation experiments using the trained nuSim to evaluate sometimes conflicting hypotheses based on their predictions about dynamical mechanisms shaping the evolution of population trajectories.

Little is known about how sensory feedback shapes the MC dynamics during movement generation. We take a modeling perspective using the developed computational framework to probe this question. Goal-driven RNN models of MC when trained to transform task-related inputs into experimentally observed EMG signals exhibit rotational dynamics [13, 20]. Therefore, it is assumed that the intrinsic recurrent connections (autonomous dynamics) of MC give rise to its experimentally observed rotational dynamics with negligible contribution from sensory feedback. However, recent studies show that models of the MC based on feedforward networks without any recurrent connections also exhibit rotational activity patterns when trained to transform proprioceptive feedback into muscle excitations, suggesting that sensory feedback contributes substantially to MC rotational dynamics [19]. We leverage the robustness of nuSim to perturbations to evaluate these conflicting theories about the role of sensory feedback in MC dynamics (Fig. 6a).

**Fig. 6.**
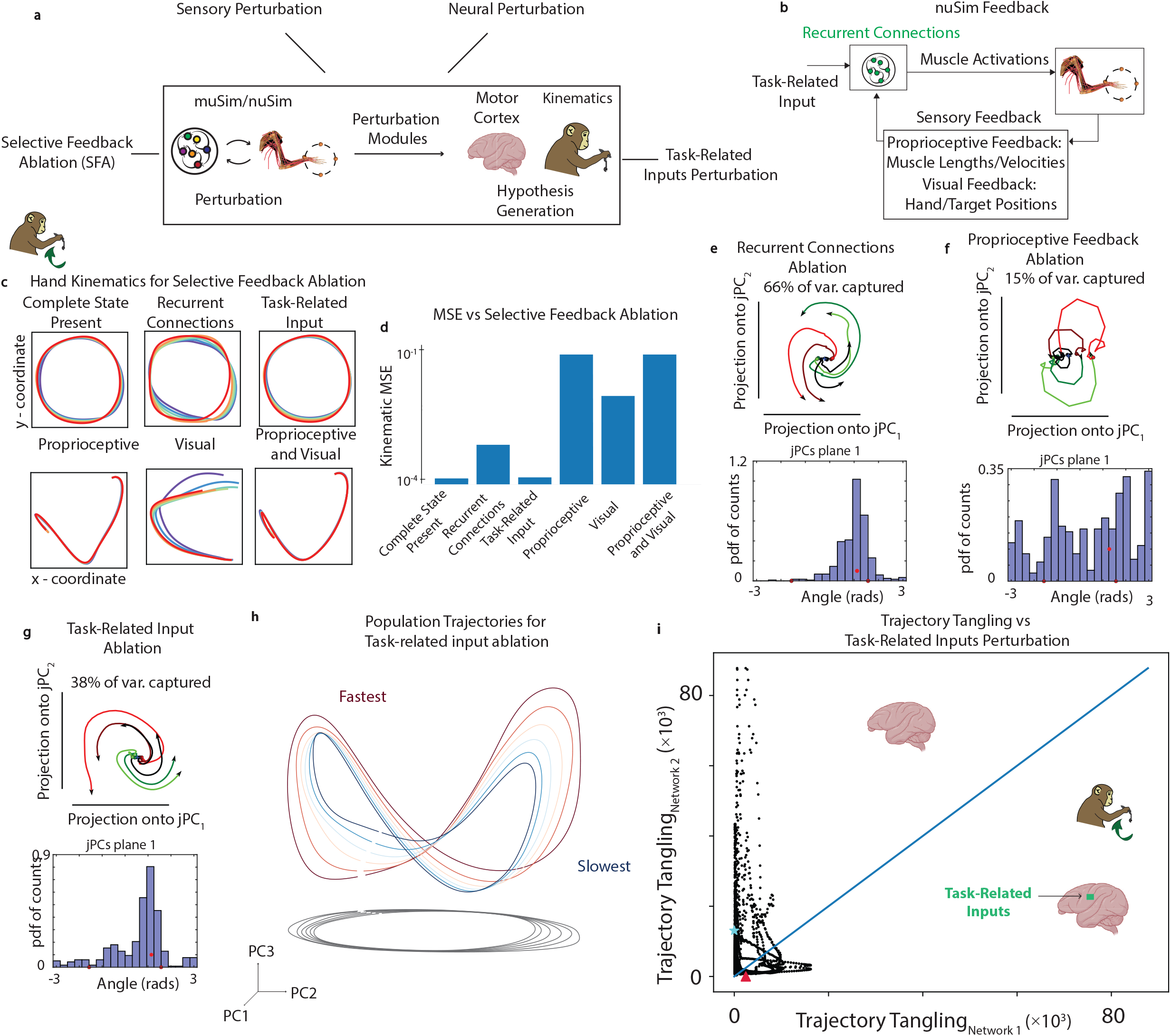
nuSim enables modeling the contribution of sensory feedback and task-related inputs towards neural dynamics and task execution. **a**. For the trained nuSim controller, various perturbation modules, such as selective feedback ablation (SFA), may allow for hypothesis generation in the neural and kinematic subspaces. **b**. Overview of the sensorimotor loop modeled using nuSim emphasizing sensory feedback and recurrent connections. The task-related input consists of a scalar signal representing a specific speed. Proprioceptive feedback consists of muscle lengths and velocities. Visual feedback consists of hand and target positions. Recurrent connections represent the recurrent weights of the RNN. **c**. Effect of various feedback ablations on movement execution (kinematics). Different colors show the hand-trajectories (x and y coordinates) for different speed conditions. The title of each subpanel shows the specific feedback that is ablated except the first subpanel for which complete state is present. **d**. MSE vs SFA. The x-axis represents the specific feedback that is ablated except for the complete state present which signifies no ablation. The y-axis represents the MSE between the experimental and nuSim kinematics averaged over x and y coordinates for all the training and testing speed conditions. **e**. jPCA projections of nuSim state for 620 ms after movement onset for all training and testing speed conditions (different colors) for recurrent connections ablation. Histogram shows the distribution of angle between the nuSim state and its derivative for all analyzed times and conditions. **f** and **g**. Same as **e** but for proprioceptive, and task-related input ablation, respectively. **h**. nuSim RNN controller trajectories in the top 3 PCs subspace for the training and testing conditions for the monkey cycling task after task-related input ablation. The separation between the elliptical trajectories for different speed conditions (different colors) is significantly reduced as compared to when the task-related input is present. **i**. Network 1 (with task-related input present) vs Network 2 (with task-related input ablated) trajectory tangling values. Points are shown for all times for the six speed conditions. Blue line indicates the unity slope. Triangle / Star represent the 90^*th*^ percentile tangling.

We performed ablation studies during the monkey cycling task by eliminating specific inputs to nuSim and observing the corresponding effects on movement kinematics and RNN activity patterns (Fig. 6). We found that the recurrent connection ablation had negligible effect on the rotational dynamics throughout the movement (Fig. 6e). nuSim was also able to achieve the task kinematics relatively well despite recurrent connection ablation (Figs. 6c and 6d). However, if the proprioceptive or visual feedback is ablated, nuSim is not able to execute the task well (Figs. 6c and 6d) and the rotational dynamics are significantly disrupted (Fig. 6f and Supplementary Fig. 7a). Our results are consistent with previous computational approaches to study the role of proprioception during motor control [81]. Therefore, specifically for the monkey cycling task, proprioceptive feedback may be more important to sustain rotational dynamics and task execution as compared to the recurrent connections.

It has been shown that MC embeds muscle-like commands in an untangled population response, even though the muscle and sensory feedback trajectories themselves are usually highly tangled [82]. Here, we test the hypothesis that task-related (condition-specific) inputs to the MC may be contributing to the observed low-trajectory tangling. The ablation of the task-specific input significantly disrupted the rotational dynamics and the separation of nuSim trajectories across different task conditions (Figs. 6g and 6h). However, nuSim achieved the task with negligible decrease in kinematic accuracy (Figs. 6c and 6d). This resulted in significantly increased trajectory tangling values (see Methods) for the nuSim RNN trajectories (Fig. 6i). This may be explained by the increased reliance of the nuSim network on sensory feedback as compared to the task-related inputs. In contrast to standard OL RNNs [20, 21], this framework allows for disentangling the role of processed sensory feedback from that of task-related inputs, presumably from the upstream brain regions and internal models, to the MC.

Lastly, we evaluate the recurrent connection ablation during the monkey cycling task with delayed sensory feedback (Supplementary Fig. 8a). We found that muSim was able to develop a representation of non-delayed sensory feedback as indicated by the high *R*^2^ between the muSim RNN activations at time *t* and the non-delayed state *s*_*t*_ (Supplementary Fig. 8b). However, recurrent connections had negligible effect on the neural dynamics and task kinematics, similar to the cycling task. We found that recurrent connections had a stronger role during the delayed reach tasks with no representation of go-cue signal in the sensory feedback. Given that nuSim dynamics are disrupted by sensory feedback, task-related inputs, and recurrent connection ablations depending on the task, we suggest that cortical dynamics are modulated synergistically in a task-dependent manner by sensory inputs, cognitive/task-specific modules and cortical recurrent connections to achieve the desired movement. Lastly, to aid future perturbation experiments using the trained muSim/nuSim, we have provided perturbation modules.

### 2.7 Discussion

We introduce a computational framework, muSim, that can be used to understand the computational role of neural activity and dynamics for the control of movement. The modularity of this framework enables the integration of musculoskeletal models of different animal species, task specifications, training algorithms, and constraints. Our adoption of complex musculoskeletal systems into the training paradigm allows an understanding of how the MC, presumably with task-relevant inputs from other brain regions, transforms high-dimensional processed sensory feedback from the upstream sensory processing regions into muscle excitations for the control of movement. Furthermore, we provide various post-training modules to analyze the underlying structure of the resulting muSim population activity after training. Emergent structures, such as rotational dynamics and stacked elliptical population trajectories, can be validated with recorded cortical data and used for further predictions. Interestingly, we found that certain population-level structures, such as orthogonal preparatory and movement subspaces during delayed reach tasks, could only be captured using muSim as opposed to standard pattern-generating RNNs.

We also provide post-training modules that can be used to quantify the dynamical alignment and similarity of muSim activity with the recorded neural data at the population and single unit level. We observed that constraining a subset of muSim units to experimentally recorded neurons during training can significantly improve their decoding for unobserved conditions. The biologically accurate modeling of the sensorimotor loop, including high-dimensional sensory feedback and neural constraints, allows muSim to be robust to perturbations, in neural, sensory and kinematic subspaces, and better generalize to unseen task conditions compared to previous models.

Our control studies show that the muscle dynamics of the musculoskeletal models and kinematic accuracy on the task play a significant role in the emergence of realistic neural dynamics in the trained muSim networks. We also provide perturbation modules that enable perturbations to be introduced at various stages of the sensorimotor transformation, such as selective feedback ablation. This framework enables inferring the effect of such perturbations on the underlying neural dynamics or task kinematics, and to infer the dynamical mechanisms shaping the neural population activity, further enabling robust hypotheses generation and testing. Using selective feedback ablation, we find that proprioceptive feedback plays an important role in maintaining the rotational dynamics of the MC in addition to the corresponding task accuracy, whereas the task-related (condition-specific) inputs to MC contributing to its autonomous dynamics help in maintaining low-trajectory tangling across task conditions. This further stresses the importance of incorporating sensory feedback transformations in modeling motor pathways and demonstrates the general hypotheses testing, including the role of internal models and multiple brain regions in movement generation, that can be performed using the muSim framework. Future work includes investigating the generalization of such properties across variations in musculoskeletal models across species, as well as performance of these models on diverse tasks.

## 2.8 Acknowledgements

This work was supported by NIH Brain Initiative grant 1RF1DA056377-01. We are very grateful to the following researchers for making their experimental data available: Abigail Russo and Mark Churchland for the monkey cycling and reaching datasets, and Claire Warriner and Andrew Miri for the mouse alternation dataset.

## 2.9 Data Availability

All datasets used in this study are publicly available. The kinematic and neural data for the monkey cycling task is available at https://figshare.com/s/b2a0557c239a1010d8ea. The kinematic and neural data for the monkey reaching task is available at https://dandiarchive.org/dandiset/000128. The kinematic and neural data for the mouse alternation task is available at https://figshare.com/articles/dataset/Neural_and_behavioral_data_from_Warriner_et_al_2022/20310684.

## 2.10 Code Availability

Code, documentation, and a demo dataset for developers and users is available at https://github.com/saxenalab-neuro/muSim.

## 3 Methods

### 3.1 Experimental Data

#### 3.1.1 Monkey Cycling Data

Head-restrained rhesus macaques were trained to perform a cycling task while sitting in a customized chair. During the controlled experiments, macaques manipulated a pedal-like device with their right-arm while the left arm was loosely restrained. The real time horizontal and vertical hand positions were recorded. The wrist movement was restrained and the cycles were driven mainly by the movement of the elbow and shoulder. This resulted in a highly stereotyped arm movement across cycles.

The macaques rotated the pedal to track a moving target with respect to their virtual first-person location on the monitor in front of them. Juice reward was dispensed so long as they maintained their virtual position close to the displayed target.

Single electrodes driven by a hydraulic microdrive were used to make sequential neural recordings from a broader range of sites in primary motor cortex and adjacent aspect of dorsal premotor cortex. For all further analyses, these recordings were treated together as a single motor cortex population. Neural signals were amplified, filtered and manually sorted using Blackrock Microsystems Digital Hub and 128-channel Neural Signal Processor. During cycling, nearly all of the isolations made were responsive and those with low signal-to-noise ratios or insufficient trial counts were discarded. Gaussian filtering (20ms SD) was then applied to filter the spikes of each recorded neuron to produce estimated firing rate, which was then trial averaged.

The trials were divided into 8 speed bins. Kinematics, EMG and neural data were then averaged across the trials in each speed bin. In each speed bin, a high degree of training ensured stereotypical behavior across trials. Adaptive alignment procedure was applied to correct remaining slight misalignments. Aberrant trials deviating from stereotypical behavior were discarded. Kinematics, neural and EMG data were then averaged across the resulting trials in each speed after applying this adaptive alignment procedure. Further details about data preprocessing are given in [21].

#### 3.1.2 Monkey Tracking Reach Data

Neural recordings for the monkey tracking reach task were made as described previously [12, 13, 83]. Single-electrode recordings were made for the monkey J from the primary motor cortex (M1) and dorsal premotor cortex (PMd) during the maze task. Monkey J performed a center-out reaching task to the displayed targets producing straight and curved reaches. We selected relatively straight reaches to the outer-targets for training. Kinematics for the generalization trajectories were generated as the average of the kinematics of neighboring two reach trajectories.

For the maze task, fixation was enforced during the preparatory period. During the movement period, relatively straight or slightly curved reaches were made. For the muSim, as this task is modeled as a tracking task without a preparatory period, we first preprocessed the recorded neural data during the movement period to control for the absence of preparatory period. Specifically, we subtracted the neural firing rate for each neuron at the start (first timestep) of the movement period from the response of that neuron throughout the movement period.

#### 3.1.3 Mouse Alternation Data

In the alternation task, mice turn a small wheel iteratively (akin to cycling) with their right forelimb. Alternation task elicited MC dependent flexor-extensor alternation. In other words, alternating activation of triceps and biceps was elicited. Mice were head-fixed and trained to perform alternation task for a water reward while muscle activity from the forelimb was recorded. Neural activity in the left caudal forelimb area was recorded after mice had been fully trained. Kinematics, EMG and neural data were then averaged across stereotypical trials from the same session.

Further details about behavior and neural data recording and preprocessing are given in [71].

### 3.2 Musculoskeletal Models

#### 3.2.1 Musculoskeletal Model of a Macaque Limb

We build on the 3D musculoskeletal model of a macaque arm developed in [61]. The developed model consists of seven degrees of freedom (DoF) that include extension and flexion of the elbow, 3D rotation about the shoulder joint, supination and pronation of the lower forelimb, adduction/abduction and flexion/extension of the wrist. The five segments represented in the model are the hand, the torso, the radial side of the lower arm, the ulnar side of the lower arm and the upper arm. Shoulder abduction and adduction is its rotation around the x–axis and its rotation around y-axis is defined as internal and external rotation. Rotation of the shoulder about the z-axis is shoulder flexion and extension. Elbow flexion is rotation about the z-axis. An intermediate x rotation is used to define the off-axis pronation and supination axis. Rotation about the x-axis at the center of the wrist is wrist flexion and extension. It is followed by rotation around the z-axis which is wrist abduction and adduction.

The model also consists of 39 muscles. These muscles are based on the anatomical data obtained from cadaveric studies [84, 85]. The muscle properties, such as muscle/tendon length, were obtained from the literature [86].

The musculoskeletal dynamics used for developing the forward dynamic model can be represented by the following equation

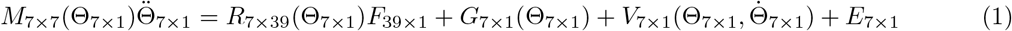

Where *M* represents the mass distribution of the system given the current joint angles Θ. 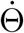 and 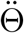 represent the joint velocities and accelerations, respectively. *R* is the moment arm matrix. *F* is a vector of muscle forces. *G, V* and *E* describe the moment contributions of gravitational, internal and external forces, respectively. The model is designed to be scalable to a generic macaque arm given the mass. Further details of the model are given in [69].

This model is adapted for use in OpenSim [87]. We first replaced the existing muscle model with a more biologically accurate and computationally stable Millard muscle model [88]. However, the forward simulations based on biologically accurate musculoskeletal models in OpenSim are computationally expensive. This makes the downstream learning of the muSim controller extremely inefficient. To overcome this challenge of slow forward simulations, we first convert the macaque arm model from OpenSim to an equivalent model in MuJoCo physics simulation engine [62, 63, 89]. MuJoCo is a state-of-the-art physics-based joint-constrained physics simulation engine that can reach forward simulation speeds of more than 600 times those of OpenSim [62]. Therefore, we optimized the OpenSim macaque arm model for use in MuJoCo by approximating the musculo-tendon units. This resulted in an anatomically accurate musculoskeletal model in MuJoCo that can facilitate the downstream learning of the muSim controller by producing fast forward simulations.

For the joint-based musculoskeletal model of macaque limb, we replaced the muscle actuators of the above anatomically accurate musculoskeletal model with motor actuators. One motor actuator was used for each DoF.

#### 3.2.2 Musculoskeletal Model of a Mouse

We utilized an anatomically accurate mouse musculoskeletal model developed in [70], and built in the PyBullet framework [90]. PyBullet is a fast, stable, and open source physics simulator with a python application programming interface (API). The authors of [70] developed a simulator-agnostic muscle library to be integrated with PyBullet for the purposes of this model. muSim controller interacts with the right forelimb of this model, directly activating its 18 muscles while the rest of the skeletal model is fixed in place. Forelimb muscle attachment points were determined based on embryonic studies of mice; due to lack of experimental data, mainly distal muscles were included. The model does not include proximal muscles that originate from the spinal segment.

For shoulder joints, which were modeled as a spherical joint, there are three rotational DoF: retraction-protraction (rotation about the transversal axis), abduction-adduction (coronal axis), and external-internal rotation (sagittal axis). For elbow joints there are two DoF: extension-flexion (transversal axis) and supination-pronation (sagittal axis). Wrist joints additionally contain two DoF: extension-flexion (transversal axis), and abduction-adduction (coronal axis). More details as well as the joint range-of-motion and limits can be found in [70].

The muscles themselves are of the Hill-type [91], with activation dynamics given by

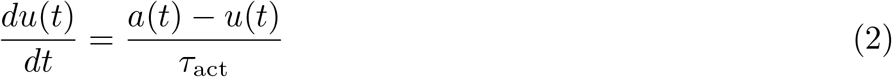

where *u*(*t*) are the activation dynamics, *a*(*t*) is the muscle excitation, and *τ*_act_ is the time constant. The muscle excitation signal *a*(*t*) is determined by muSim controller, directly controlling the muscle activation in order to produce the necessary movements. We chose a biologically realistic starting position of the forelimb based on the joint range-of-motion given in [70].

### 3.3 Kinematics MSE

The MSE between the kinematics produced by muSim and experimental kinematics is calculated as follows:

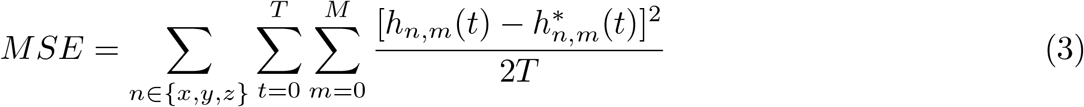

where *x, y* and *z* represent the markers’ x, y and z coordinates, respectively. *h*^∗^ and *h* represent experimental and muSim markers’ kinematics, respectively. *T* represents the total number of time points in the corresponding training condition trajectory.

### 3.4 Initial Pose Estimation and Movement Trajectory Feasibility

To estimate the initial pose for the musculoskeletal model and to determine if the user-specified movement trajectory is feasible, we first define the following objective function:

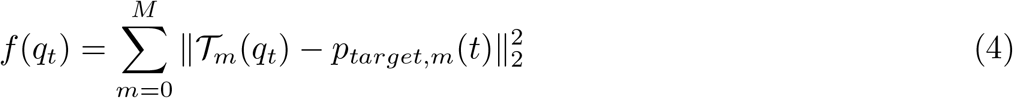

where *q*_*t*_ is a vector consisting of joint positions of the musculoskeletal model, and *p*_*target*_(*t*) consists of the user-specified marker targets positions in the cartesian coordinate plane. 𝒯_*m*_ is a function that transforms *q*(*t*) into the cartesian coordinates for the corresponding user-specified marker/target m.

To figure out the initial pose for the musculoskeletal model, we provide Trust-Region optimization algorithms (Trust Region Optimization [92]). Specifically, the following objective function is minimized:

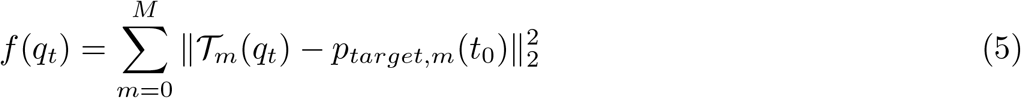

where *p*_*target,m*_(*t*_0_) represents the initial position of the target/marker *m* specified by the user.

In cases, where the straight path trajectory from the default pose of the musculoskeletal model to the user specified initial targets’ positions is not possible, we provide the evolutionary algorithms (covariance matrix adaptation evolutionary strategy (CMA ES) [93, 94]) to optimize the objective function defined in (5).

To figure out if the movement trajectory as specified by the user-specified kinematics is feasible for the musculoskeletal model, we provide optimization algorithms to minimize the following objective function:

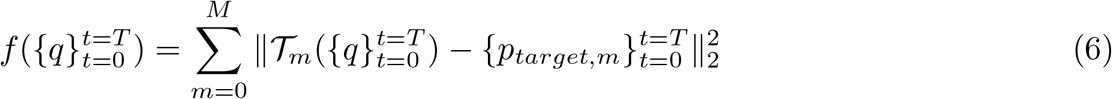

where, 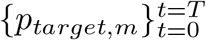 represents the target *m* trajectory specified by the user and 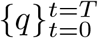 represents the trajectory of joint positions as generated by the optimization algorithm.

### 3.5 Formulation of the Reinforcement Learning Problem

We formulate the task of controlling musculoskeletal models to perform different movements as a reinforcement learning problem. This formulation consists of a policy network, critic networks and an environment. Given the current state *s*_*t*_ ∈ *S* of the environment, the policy network outputs the probability distribution, represented by *π*_*θ*_(*a* | *s*), over the possible actions *a* ∈ *A. θ* denotes the parameters of the policy network implemented using RNNs. The environment can be implemented using different physics simulation engines. In this work, we use MuJoCo [89] and PyBullet [90] physics engines for implementing the environment. The environment consists of musculoskeletal models that can interact with other physically simulated objects. Advanced physics engines allow the implementation of physically realistic simulations under different constraints such as contacts. At each timestep *t*, the environment receives an action *a*_*t*_ from the policy network. *a*_*t*_ represents the control input or motor command. Given *a*_*t*_, the esulting movement is executed and the environment transitions from the current state *s*_*t*_ to next state *s*_*t*+1_. *s*_*t*+1_ represents the physical effects of the motor command executed in our biomechanical simulation. The state transitions can be stochastic in muSim and the conditional probability of reaching the next state *s*_*t*+1_ is given by *p*(*s*_*t*+1_| *s*_*t*_, *a*_*t*_). The state transitions are governed by the underlying dynamics of the musculoskeletal system and the physics environment. These dynamics are defined by differential equations represented by *F* :

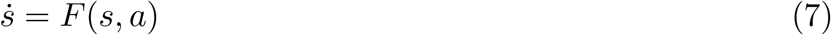

Given the probability of starting in some initial state *s*_0_, the probability of realizing a trajectory *T* = (*s*_0_, *a*_0_,, *s*_*N*_ ) under the policy *π*_*θ*_(*a*|*s*) is given by:

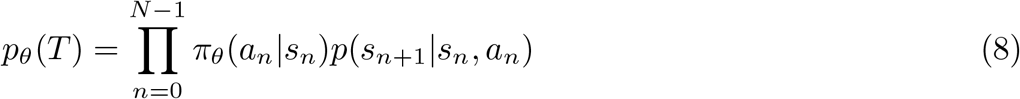

In current work, we implement deterministic transitions of the environment. At each timestep *t*, a reward *r*_*t*_ is also generated to quantify the performance of the policy network.

### 3.6 Environment Design

The environment can consist of anatomically accurate musculoskeletal models of different animal species that can interact with other objects while obeying physical laws and constraints, such as collisions, friction, torque, contacts and ground reaction forces. This can allow the simulation of freely moving animal species that can interact with their environment.

In current work, the objective is to reproduce experimentally recorded movement. Some portions of the musculoskeletal models in this work are fixed in space while the limbs/extremities and thus the end-effector can move freely. The experimentally recorded kinematics are represented in the simulation using abstract objects known as targets. At each timestep *t*, the position of the target is updated to reflect the experimentally recorded position to be tracked by the end-effector in a 3D plane. The performance of the policy network at each timestep *t* can thus be quantified by the distance between the end-effector position and the target position. The policy network is then trained to output muscle commands such that the end-effector tracks target position at each timestep *t*, thus reproducing the experimentally recorded behavior.

The same framework can be used to model the task or abstract behavior in the absence of experimentally recorded kinematics using sparse metrics for quantifying the performance of the policy network. For example, in the absence of experimental kinematics, the reaching task can be achieved by fixing the abstract target at the desired final position, such as in the monkey delayed reach experiments.

### 3.7 States/Actions

The environment state s_*e*_(*t*) is filtered to form the state feedback *s*_*t*_ for the policy network. *s*_*t*_ may consist of complete or partial information required to solve the control problem. If partial information about the environment state s_*e*_(*t*) is available such that the state feedback is imperfect or partially observable, the control problem in the DRL framework can be formulated as a partially observable Markov decision process (POMDP). Various formulations of DRL, such as deep hierarchical RL, can be used to solve the control problem for POMDPs [95]. In previous work, a DRL formulation with RNN implementation of the policy network has also been shown to solve control problems with a partially observed state [96]. This also justifies the use of RNN implementation of the policy network in current work, in addition to the existence of strong recurrent connections in the motor pathways. This may also point to the fact that the motor pathways may be able to construct complete state from partially observed information through these recurrent connections. In a dynamic environment with freely moving subjects, often only partial state information is available. Therefore, this computational framework can be used to solve such complex tasks.

In this work, *s*_*t*_ represents the sensory feedback and consists of both proprioceptive and visual feedback. Specifically, state feedback *s*_*t*_ to the policy network consists of muscle lengths and velocities (proprioceptive feedback) and a difference vector representing the distance between the end-effector and the target, and a condition-specific scalar representing different conditions within a task (‘task information’). The task/condition-specific scalar was included to model any input to the MC that may be contributing towards its autonomous dynamics.

The action vector *a* ∈ *A* represents the control inputs or motor commands that are then transformed into muscle activations. These activations actuate some DoF by applying motor torque. Physics simulation engines, such as MuJoCo, can also accurately model additional active and passive forces, including gravitational and contact forces, in addition to these actuated forces. Such additional forces are not modeled accurately using biomechanics engines, such as OpenSim. Therefore, such biomechanics engines can not be used to model neural dynamics accurately in the context of musculoskeletal models interacting with each other or other objects in the environment. The developed computational framework is therefore expected to advance our understanding of motor control in such complex tasks.

### 3.8 Reward Function Design

The reward function *r*(*s*_*t*_, *a*_*t*_, *n*_*t*_) specifies the behavior of the policy network. The reward function consists of two parts: task specification *T* (*s*_*t*_, *a*_*t*_) and constraints specification *C*(*s*_*t*_, *a*_*t*_, *n*_*t*_).

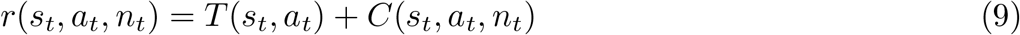

where, *n*_*t*_ represents neural parameters, such as synaptic weights or firing rates. In the developed computational framework, these neural parameters *n*_*t*_ correspond to the parameters *θ* and activity 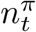 of the policy network. To keep the notation concise, we assume *n*_*t*_ ⊂ *s*_*t*_.

The generated rewards can be immediate, such as provided at each timestep *t*, or they can sparse, such as provided only at the last timestep.

It has been proposed that the central nervous system (CNS) achieves a specific task under various kinematics- and behavioral-level constraints. Previously, constraints based on the minimization of muscle effort [97, 98] or maximizing the smoothness of the end-effector trajectory and of the torque commands have been proposed [99, 100]. The models based on these constraints have been successful at reproducing empirical kinematic data. However, it is not clear how these constraints effect the CNS. Other models assume noise in the motor command that scales with its magnitude [101]. The CNS aims to then minimize the end-point variance given this motor noise.

However, is also not clear if such scaling is a consequence of neural constraints, such as neuronal noise. Therefore, it remains an open question how we approach this constraint for novel, unrehearsed movements [98].

In this work, we instead assume and validate the existence of neural constraints, such as minimization of neural firing rates, that are suitable from an energy minimization and evolutionary standpoint. These neural constraints can also be thought of giving rise to other previously proposed kinematics- and behavioral-level constraints. We also validate that such constraints generalize across novel and unrehearsed movements. To test specific hypotheses, this framework can be used to implement various behavioral-and neural-level constraints, such as the ones described above. After training, the resulting behavior and network activity under specified constraints can then be analyzed or compared with experimental data to validate their existence.

For the purposes of this work, these neural constraints can be implemented using regularizations on the policy network and are described below. Here, we design the reward function as consisting purely of the task specification *T* (*s*_*t*_, *a*_*t*_). In this work, for the tracking tasks, the task specification is designed to make the end-effector reproduce the experimental kinematics by tracking the target at each timestep *t*:

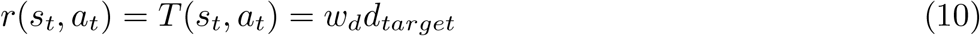

Where

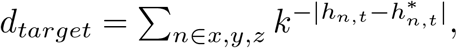

*h*_*x,t*_, *h*_*y,t*_ and *h h*_*z,t*_ represent the x, y and z coordinates of the end-effector’s position and 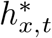, 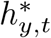 and 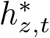 represent the x, y and z coordinates of the target’s position at timestep *t*. This reward function *r*(*s*_*t*_, *a*_*t*_) produces the maximum reward *r*_*t*_ for the motor command that results in the minimum distance between the end-effector and the target for the given timestep *t. r*_*t*_ is designed to decay exponentially with the increasing distance between the end-effector and the corresponding target to accelerate the learning of the policy network parameters. We use *w*_*d*_ = 5.0 for the monkey cycling task.

For the monkey delayed reach task, we set 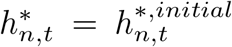 during the preparatory period, with 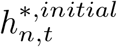 representing the initial position of the target or fixation point during the preparatory period. During the movement period, we set 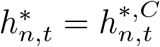 which represents the final position of the target for the reach condition *C*.

In the muSim framework, we provide two types of DRL-based training paradigms: 1. Tracking-based training, and 2. Task-based training. For tracking-based training, an immediate reward is provided at each timestep using the specified experimental kinematics. For task-based training, experimental kinematics are not required and the task-specification can be more abstract. However, the training can be challenging due to sparse rewards being provided. To overcome this challenge, we have used curriculum learning. In curriculum learning, the muSim controller is trained in two stages. In the first stage, the controller is trained to gradually build up an increasingly complex repertoire of behaviors using tracking-based training. In the second stage, the controller is further trained using the task-based paradigm. This curriculum learning can also be adjusted to match the training of animals in experiments.

### 3.9 Soft Actor-Critic (SAC) Algorithms with added neural regularizations

We adapt the SAC algorithm to train the policy and critic networks to achieve the desired task. SAC is an off-policy RL algorithm based on the maximum entropy framework [102]. This algorithm is chosen because of the complex environmental dynamics and computational complexity of the biomechanical simulations. The additional entropy regularization aids in exploration for such cases.

The reward *r*_*t*_ generated at each timestep *t* determines the behavior of the policy trained using this algorithm under the neural constraints. The sum of the rewards discounted by the factor *γ* defines the return R.

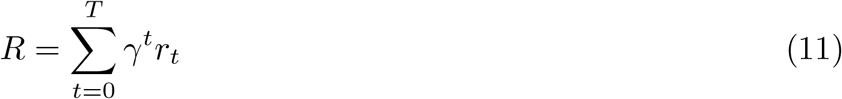

Where *T* is the last timestep. The discount factor *γ <* 1 determines the relative importance of the future rewards relative to the earlier rewards. We set *γ* = 0.99 for training muSim.

The high dimensional action space and the complex environmental dynamics in biomechanical simulations make exploration quite inefficient using standard RL algorithms. Here, an entropy term *H*(*π*_*θ*_(.|*s*)) is incorporated in the return to achieve efficient exploration required for training.

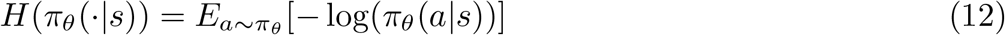

The entropy term determines the stochasticity of the trained policy and is used to achieve efficient exploration and robust convergence towards the global optimum given the high dimensional action space and complex environmental dynamics. Probabilistic matching that has been used to explain human decision making can be considered the biological analog of the maximum entropy RL framework [103].

The objective is to learn a policy that maximizes the following soft return:

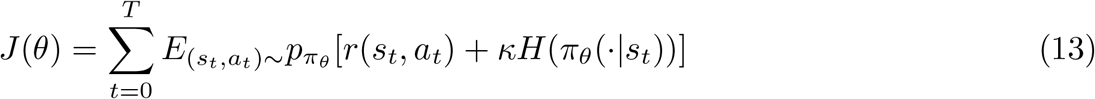

The temperature coefficient *κ* controls the stochasticity of the trained policy by determining the relative importance of reward against the entropy term. *κ* is adjusted automatically during training using the dual gradient-descent implemented in the SAC algorithm [102].

The SAC algorithm uses policy iteration to train the parameters of the policy network to maximize the objective function *J*(*θ*).

### 3.10 Value Network

The value/critic network represents the transformation from the state-action pair (*s*_*t*_, *a*_*t*_) to its q-value *q*(*s*_*t*_, *a*_*t*_) which in turn represents the expected soft-return of motor command *a*_*t*_ given the sensory feedback *s*_*t*_ in the maximum entropy RL framework [102]. The critic network parameterizes the q-value *q*_*ϕ*_ through parameters *ϕ*. In this work, the critic network consists of three feedforward layers with 256 nodes each.

Dopaminergic projections from the ventral tegmental area to the MC may reflect the neural correlates of reward signals (or reward prediction errors) [104]. Regions of the brain involved in reward processing, such as basal ganglia along with these dopaminergic projections, can be considered analogous to the critic network [105].

### 3.11 Policy Network

The actor/policy network parameterizes the policy *π*_*θ*_ through the parameters *θ*. The policy network represents the mapping from state feedback *s*_*t*_ to the probability distribution over the control inputs space. In the brain itself, various regions are involved in the transformation from the sensory feedback to the motor command. The sensory feedback is first processed in the sensory and visual processing regions of the brain that share strong reciprocal connections with the premotor areas. The premotor cortex processes the feedback further and has strong reciprocal connections to the MC. The MC then transforms the processed feedback into motor commands through subcortical and spinal cord projections [22, 59].

We base the architecture of the policy network to mimic the modular structure and reciprocal/recurrent connections of the motor pathways involved in the movement generation. The policy network consists of three layers representing the sensory (input) and premotor/motor regions (recurrent) and the final (output) subcortical and spinal cord projections. The layers representing the premotor and motor regions are based on RNNs to mimic the recurrent connections. For comparison with the recorded cortical data, we use the activity of the RNN layers representing the premotor and motor cortex.

The policy network consists of three layers. The first layer is a feedforward forward layer with the following input-output transformation:

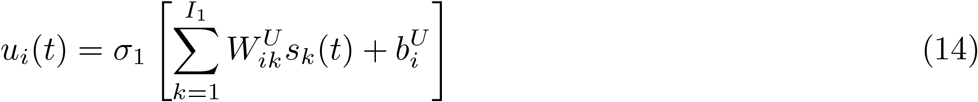

where *s*_*k*_(*t*) is the input sensory feedback with dimensionality *I*_1_. *σ*_1_ is the non-linearity for the first layer. *W*^*U*^ and *b*^*U*^ represent the weights and biases for the first layer.

The second layer consists of an RNN. RNN can be considered a discrete dynamical system with the following dynamics:

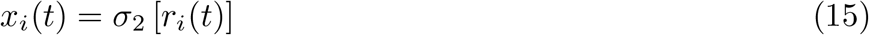

with

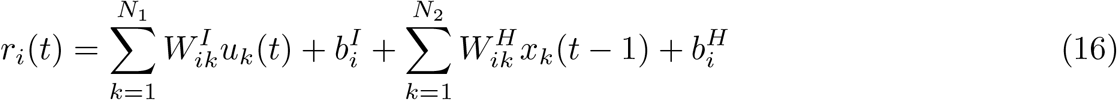

Where *r* represents the inputs to the non-linear activation function *σ*_2_ and *x* represents the corresponding output of RNN hidden layer. *N*_1_ is the number of units in the first feedforward layer and *N*_2_ is the dimensionality of the hidden layer of the RNN. *σ*_2_ is the non-linearity for the RNN layer. *W*^*I*^ and *b*^*I*^ represent the input weights and biases, respectively. *W*^*H*^ and *b*^*H*^ represent the recurrent weights and biases for the RNN layer, respectively.

The final layer is a feedforward layer with the following input-output transformation:

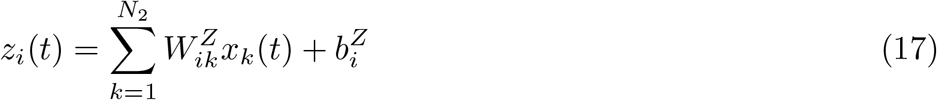

The output of the third layer, *z*, represents the muscle command with dimensionality *N*_3_. *W*^*Z*^ and *b*^*Z*^ represent the weights and biases for the final feedforward layer, respectively.

### 3.12 Neural Regularizations

Here, we hypothesize that the brain is evolved to produce optimal behavior under neural constraints. These neural constraints can be implemented as regularizations on the networks. Here, we implement three neural constraints as regularizations on the policy network activity.

The first regularization term is an *L*_2_ penalty on the input and output weights of the three layers which encourages the minimization of synaptic weights (SW) or sparse connections between the network nodes.

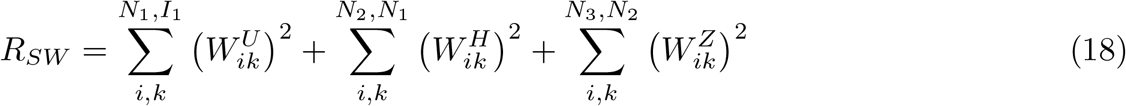

The second regularization term encourages the minimization of the neural firing rates (FW). It also prevents the network activity from saturating:

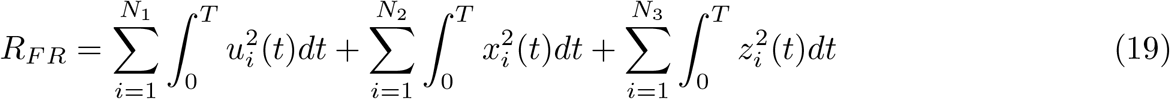

Where *T* consists of cumulative timepoints for all the conditions the network is trained on.

Finally, we implement a third regularization term that encourages the network to achieve the task while making simple state-space trajectories (SD, simple dynamics) as proposed in [20].

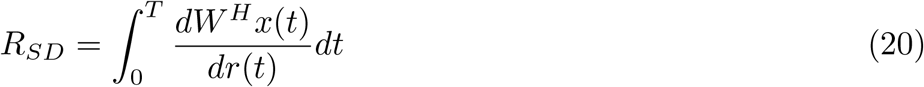

Therefore, the final loss function consists of the following terms:

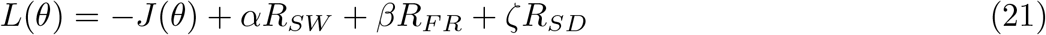

Where *θ* = *W*^*U*^, *W*^*H*^, *W*^*I*^, *W*^*Z*^, *b*^*U*^, *b*^*H*^, *b*^*I*^, *b*^*Z*^ represents all the parameters of the policy network.

The policy network parameters are then trained using gradient descent to minimize the loss *L*(*θ*). We use Adam optimizer to update the parameters of the loss function. This loss function is used for muSim training.

We used *α* = 0.001, *β* = 0.01 and *ζ* = 0.1. We used *N*_1_ = *N*_2_ = 256. Additionally, we used *σ*_1_ = *σ*_2_ = *tanh*.

### 3.13 Simultaneous goal- and data-driven modeling

To implement the simultaneous goal- and data-driven modeling (GDM), we enforce an additional constraint on a subset of the RNN units to follow recorded neural activity for the training conditions. Specifically, we minimize the following loss between the network and recorded single unit activity:

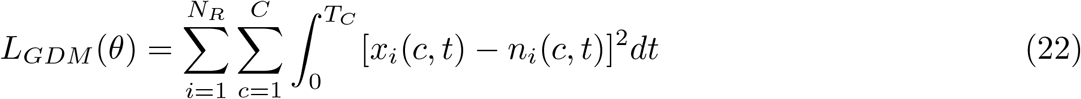

Where *T*_*c*_ represents the number of timesteps per condition, *C* is the total number of training conditions, *n* represents recorded neural activity, and *N*_*R*_ is the total number of recorded neurons. At each timestep *t*, the simultaneous goal- and data-driven modeling loss minimizes the difference between the activities of the subset of nuSim controller’s RNN units and the corresponding recorded neurons. The final loss is:

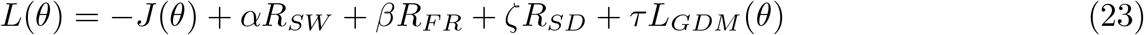

This loss function is used for nuSim training.

We used *τ* = 10^4^ to account for relatively less number of experimentally recorded neurons as compared to total number of units in the policy network and relatively smaller magnitude of *L*_*GDM*_ as compared to other terms in (23).

### 3.14 CCA

We used CCA to find the correlations *R*^2^ between the trained muSim network and the experimentally recorded neural population responses [106]. CCA finds the weightings for the units in network and experimental datasets such that the reweighted datasets are maximally correlated. Both the network and recorded neural activities were first reduced to ten dimensions using PCA. CCA was then applied to recover the subspace of maximum correlations between the recorded and network activities. Inverse CCA was applied to transform the muSim network activities from the maximum correlations subspace back into the recorded activities subspace. We reported the reconstruction comparison and *R*^2^ between the network and experimentally recorded activities in this subspace.

### 3.15 jPCA

We used jPCA [13] to quantify the oscillatory dynamics in neural state *x*(*t, c*) across times, *t*, and conditions, *c*. jPCA provides summary features, such as quality of fit and variance explained, relevant to the hypothesis that the neural state evolves according to the oscillatory dynamics. It also allows the visualization of the two-dimensional projection of the neural data containing the oscillatory dynamics.

### 3.16 Linear Regression Analysis

We used linear regression analysis (LRA) to compare the muSim network and experimentally recorded single-unit-level neural activities. Using LRA, we first fit a linear model with ridge regression on all the conditions except the held-out condition. The network and neural activities for training conditions excluding the held-out condition are first concatenated along the time dimension separately. Ridge regression is then used to fit a linear model in which the concatenated activity for each recorded neuron is determined by the concatenated muSim network activity separately. For testing, we used this trained linear model to transform the network activity for each held out condition into the recorded neural activity subspace. The transformed network activity is then compared with the corresponding recorded single-unit level neural activity for the held-out condition to find the correlations. We used a similar procedure for comparing the kinematics and EMG models.

We used a regularization coefficient of 0.05 for LRA.

### 3.17 Procrustes Analysis

We used Procrustes Analysis to compare the network and recorded neural single-unit level activities [78, 79]. Procrustes applies linear transformations, such as scaling and rotation, to the network activity to align it with the neural activity for the given condition. Procrustes minimizes the following loss represented using the disparity (D) measure:

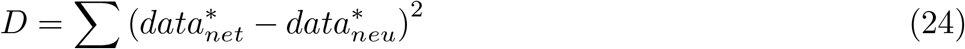

where 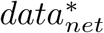 and 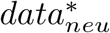 represent the muSim network activity after linear transformations by Procrustes and the standardized recorded neural activity, respectively. The rows of the data matrix correspond to the timepoints and the columns correspond to the units/neurons. We first apply PCA to the two activities to reduce the dimensions to the lesser of the two activities.

### 3.18 Trajectory Tangling

Trajectory tangling was quantified using

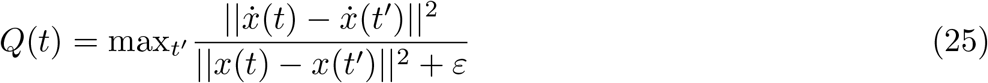

where *x*(*t*) is the muSim/nuSim RNN’s state in the top 3 PCs subspace at time *t*. 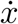 represents its temporal derivative and was calculated using 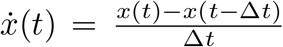 with Δ*t* = 1ms. ||.|| represents the Euclidean norm and *ε* was set to 10^−3^ to prevent the division by 0.

## 4 Supplementary Figures

**Supplementary Fig. 1.**
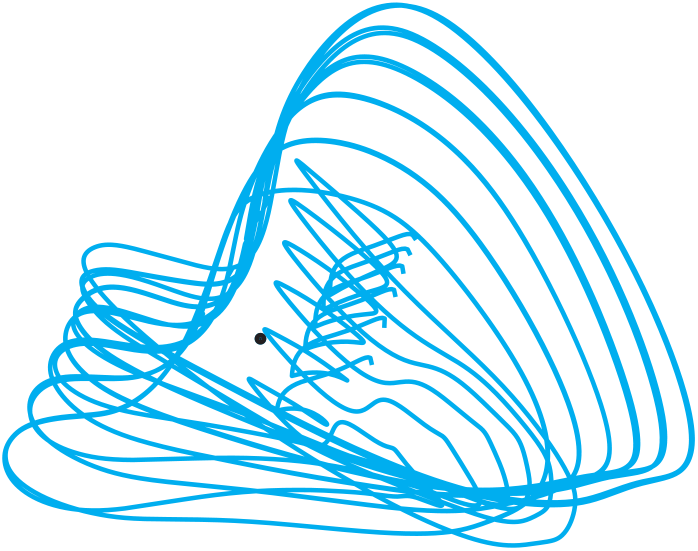
muSim RNN exhibits condition-independent fixed-point structure for the monkey cycling task. Fixed point analysis identified a single stable fixed point (black dot) for the muSim RNN trained for the monkey cycling task. muSim RNN trajectories for the different speed conditions (cyan) traverse about this stable fixed-point in a stacked elliptical structure. All the trajectories are visualized in the 3-dimensional subspace determined by the top 3 PCs computed across the six seen and unseen speed conditions. The lack of rich fixed-point structure for the monkey cycling task, as opposed to the monkey delayed reaching task, suggests that the muSim controller may be mainly sensory feedback driven as opposed to internal recurrent dynamics.

**Supplementary Fig. 2.**
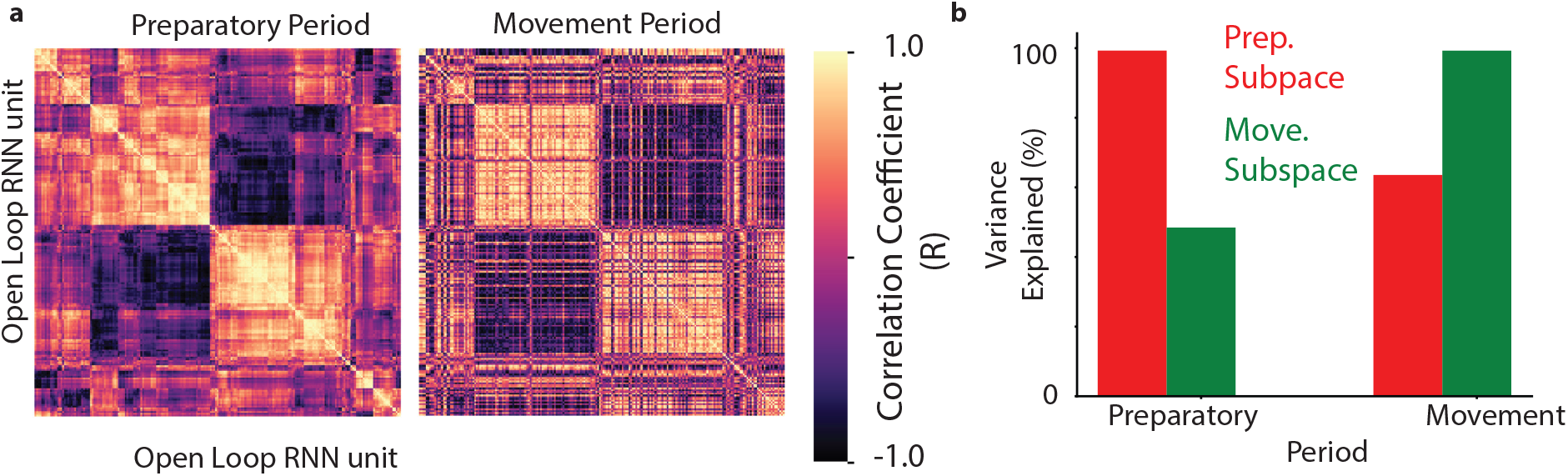
Open Loop (OL) RNN trained to transform sensory feedback into muscle excitations obtained from the trained muSim does not develop orthogonal subspaces. **a**. Preparatory period (left) and movement period (right) correlation matrices for the OL RNN units. Each entry in the matrix represents the degree to which the response was similar for the two units during that period. **b**. Percentage of variance captured by the preparatory (red) and movement (green) subspaces during the two epochs. The muSim RNN response during each epoch was projected onto the top ten dimensional preparatory and movement subspaces. The left pair of bars corresponds to the variance captured during the preparatory period. The right pair of bars corresponds to the variance captured during the movement period.

**Supplementary Fig. 3.**
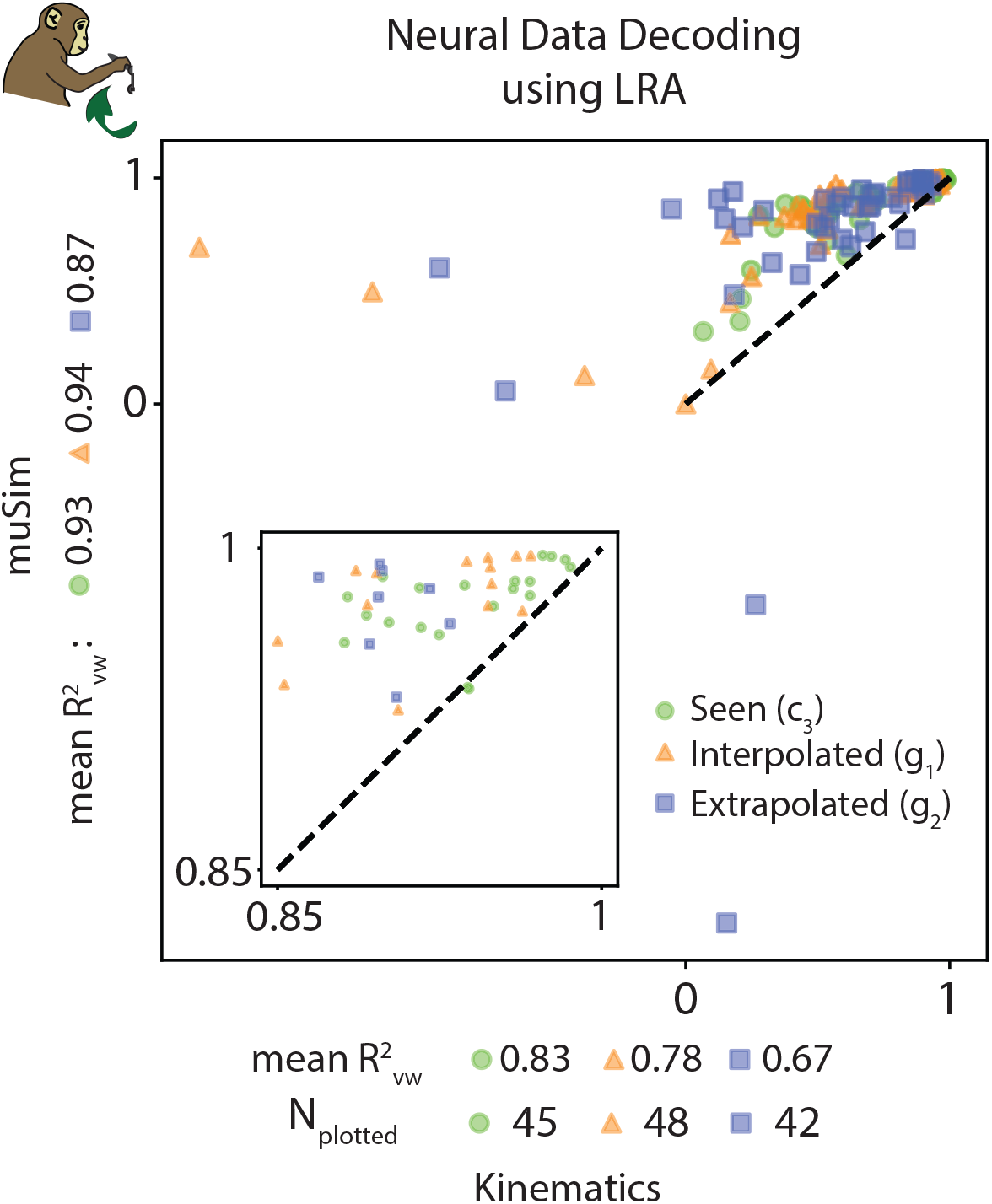
Comparison of muSim single-neuron decoding accuracy against that of kinematics-based representational model of MC. Scatter plot shows the comparison of LRA-based single-neuron decoding accuracy (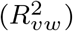) of muSim against the kinematics-based representational model of MC on the seen, interpolated and extrapolated speeds. For each neuron (dot), the *R*^2^ of muSim RNN is shown along the vertical axis and *R*^2^ of the alternative model is shown along the horizontal axis. If a neuron lies above the dashed unity line, it is better predicted by the muSim as compared to the alternative model. Only those neurons are shown for which the *R*^2^ ≥ 0 for either of the two models. The total number of neurons plotted *N*_*plotted*_ for each condition are also indicated. The variance-weighted mean 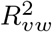 values between the experimentally recorded firing rates and their reconstructions are also shown. Inset shows the zoomed in version of the outer scatter plot.

**Supplementary Fig. 4.**
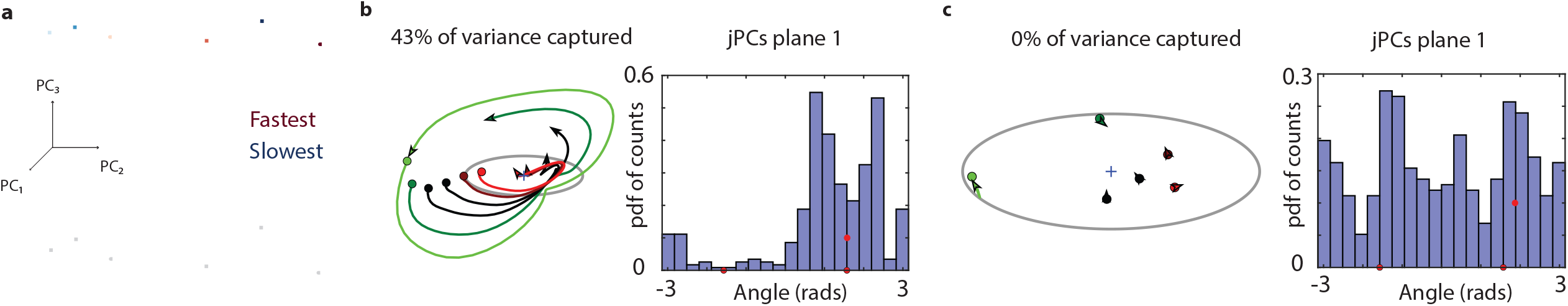
jPCA shows that the joint-based and random-muSim have distorted rotational dynamics as compared to muscle-based muSim. **a**. Randomly initialized muSim RNN trajectories for different speed conditions (different colors) for the monkey cycling task. muSim RNN trajectories settle at speed specific fixed-points after transient initial activity. **b**. jPCA trajectories for the seen and unseen speed conditions (different colors) for the monkey cycling task for the joint-based muSim. Each trace (left panel) shows the evolution of the muSim state over 620 ms after movement onset for a given speed condition. The variance captured by the jPCs plane reduces slightly as compared to the muscle-based muSim. However, state trajectories now show a bimodal angle distribution, with an increased contribution from angles greater than 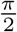 (right panel). **c**. Same as **b** but for the random muSim. The state trajectories quickly settle at speed specific fixed-points after the initial transient activity and show no rotational dynamics.

**Supplementary Fig. 5.**
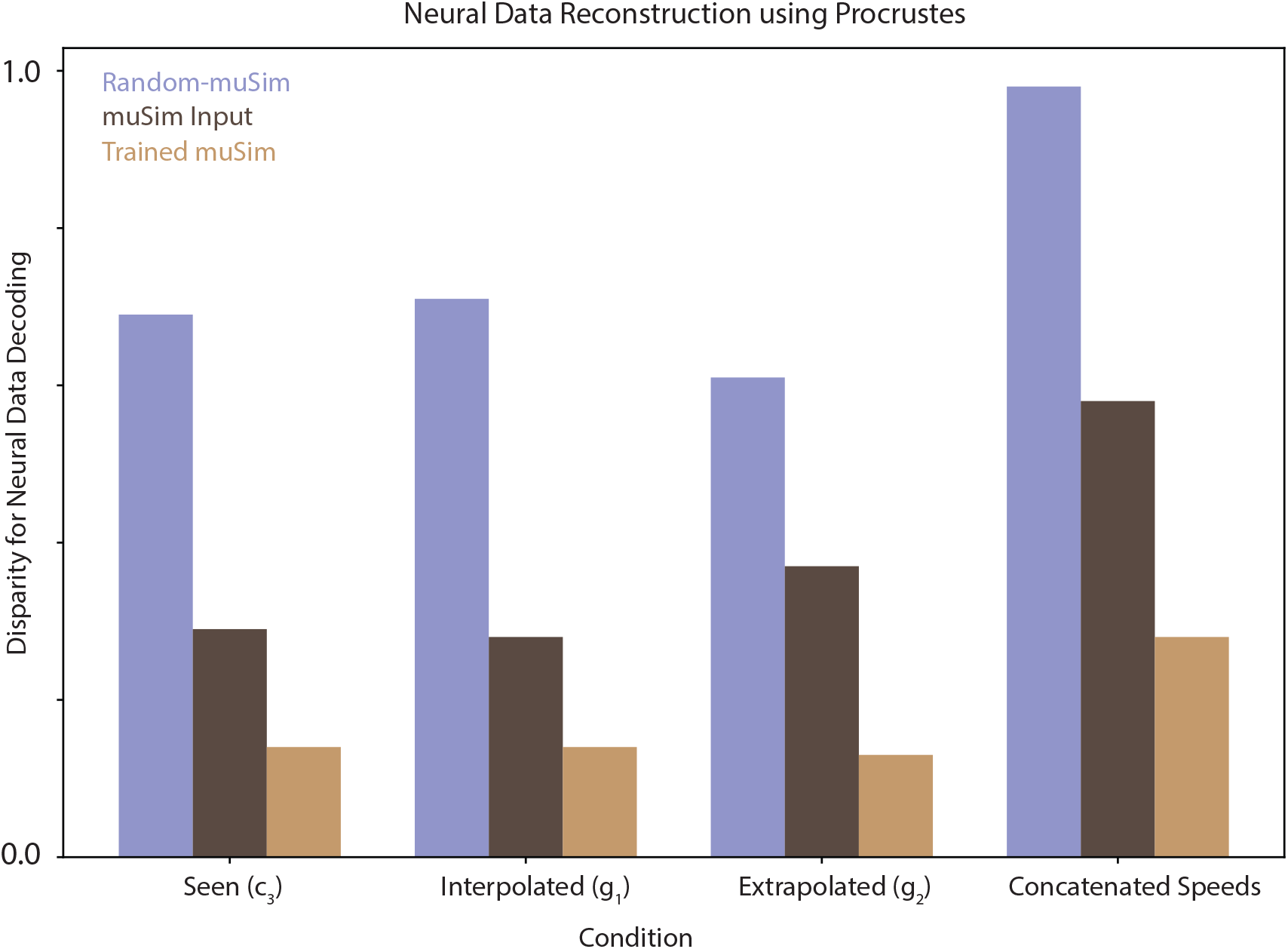
Trained muSim shows the highest alignment with the recorded neural data as compared to the random-muSim and muSim inputs. We used Procrustes to compare the recorded neural data decoding accuracies of trained muSim (red bar), muSim inputs (brown bar) and random muSim (blue bar) for the seen, interpolated, extrapolated and concatenated speed conditions. Trained muSim showed the lowest disparity followed by the muSim inputs.

**Supplementary Fig. 6.**
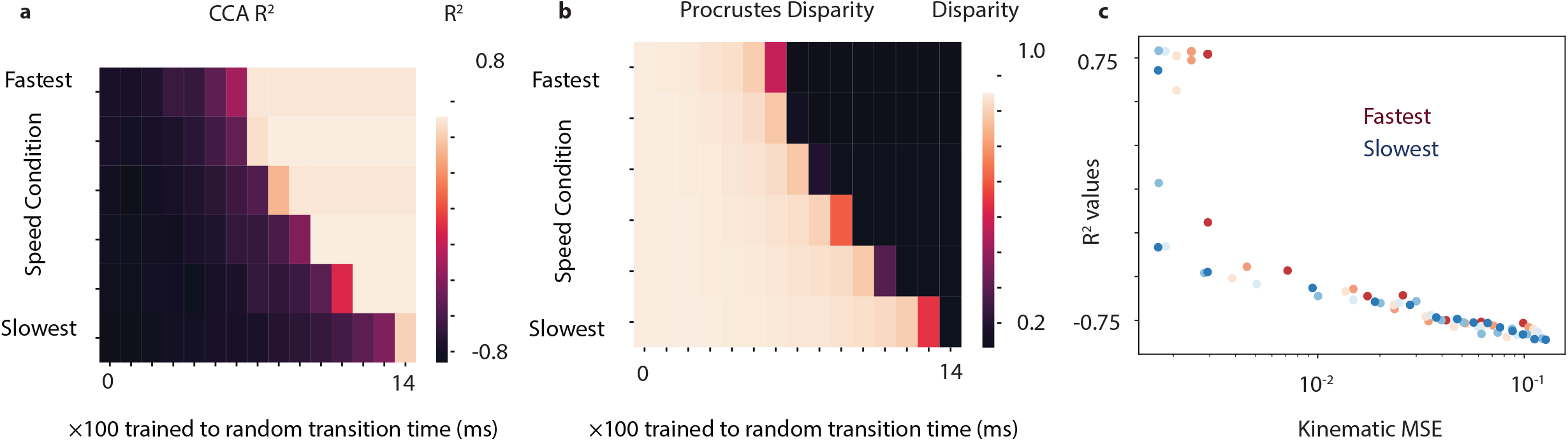
Control studies show that kinematic accuracy of muSim has exponentially increasing relation with CCA-based correlation values and exponentially decreasing relation with disparity values. **a**. Heatmap comparing the CCA-based *R*^2^ values for all speed conditions against the trained to random transition time. The trained to random transition time represents the time when the trained muSim RNN is replaced with randomly initialized RNN. **b**. Heatmap comparing the procrustes-based disparity values for all speed conditions against the trained to random transition time. **c**. Scatter plot comparing the CCA-based *R*^2^ values against the kinematic MSE for the seen and unseen speed conditions (different colors). Lower kinematic MSE results in exponentially higher *R*^2^ values suggesting a higher alignment between the muSim and experimentally recorded neural states for all speed conditions.

**Supplementary Fig. 7.**
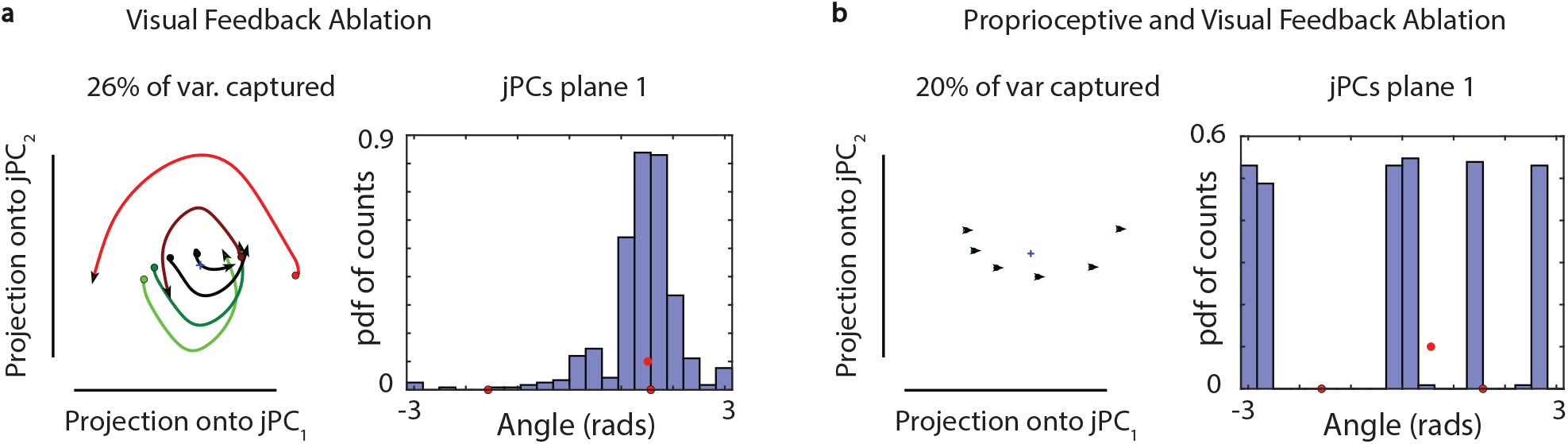
Modeling the contribution of sensory feedback towards neural dynamics using muSim. **a**. jPCA projections of nuSim state for 620 ms after movement onset for all training and testing speed conditions for the visual feedback ablation. Histogram shows the distribution of angles between the nuSim state and its derivative for all analyzed times and conditions. **b**. Same as **a** but for the proprioceptive and visual feedback removal.

**Supplementary Fig. 8.**
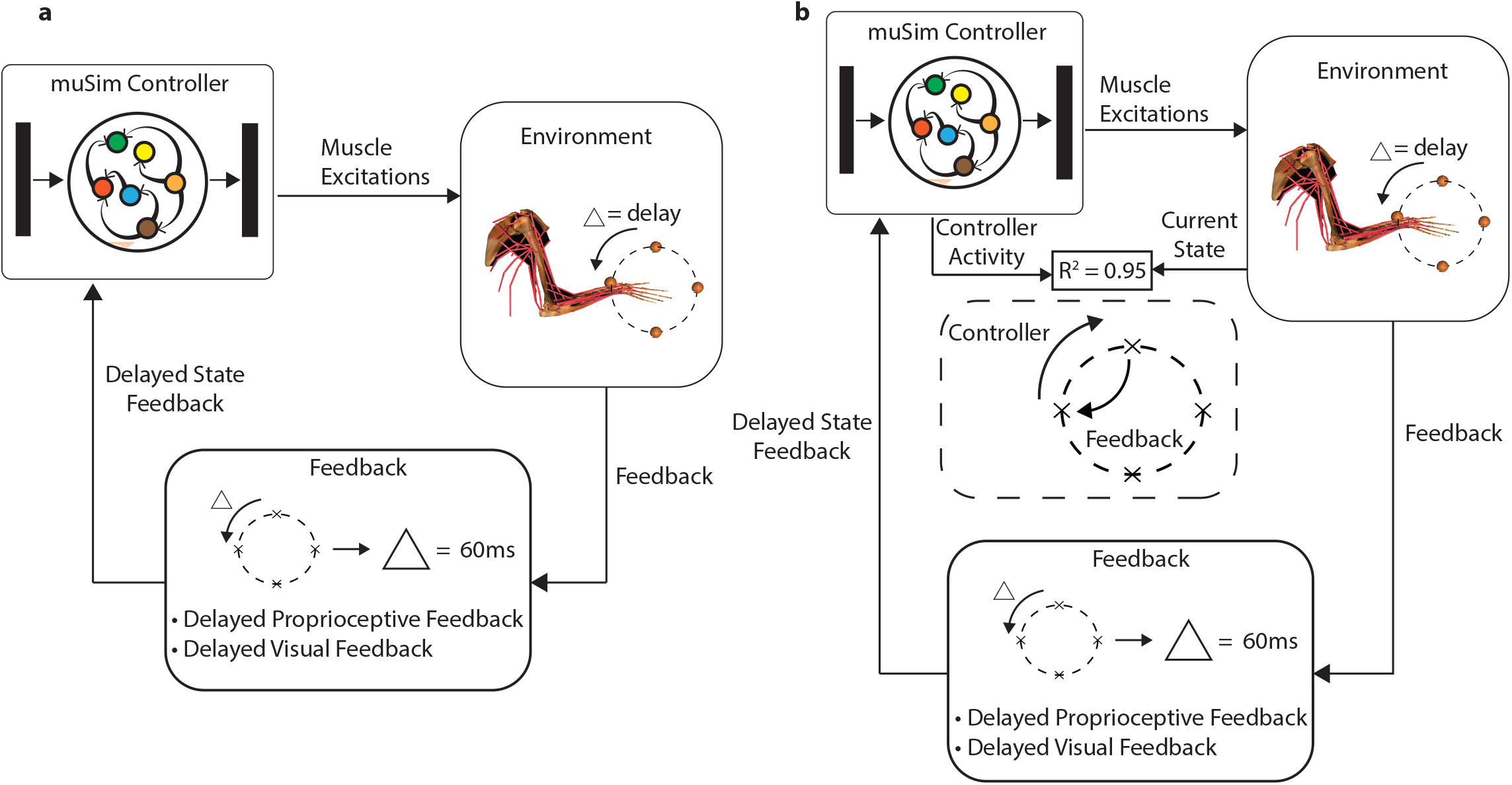
muSim can model the sensorimotor feedback delays. **a**. Schematic showing the goal-driven sensorimotor framework, muSim, in the presence of feedback delays. The current musculoskeletal and environmental state is delayed by 60ms constituting the delayed sensory feedback. This delayed feedback is then fed to the muSim controller as input. **b**. For the muSim controller trained with delayed state feedback, we observe a high mean *R*^2^ between the muSim controller’s activity and current state feedback. The feedback is delayed by 60 ms but the muSim may develop a representation of the current state to solve the task.

